# HOT1 functions as a homodimer to promote ALT-permissive chromatin

**DOI:** 10.64898/2026.11.02.763659

**Authors:** Alexia Hillairet, Nurkaiyisah Zaal Anuar, Nivedita Kumar, Vartika Khanchandani, Jia Hui Tee, Michelle Yaochai, Chai Yeen Goh, Falk Butter, Lars Nordenskiöld, Aghil Soman, Dennis Kappei

## Abstract

Telomere homeostasis is a tightly regulated and dynamic process sustained by a network of proteins that organize telomeric chromatin and safeguard chromosome stability. In ALT-positive cells, telomere maintenance relies on recombination-based mechanisms and is modulated by local chromatin states and epigenetic regulation. Here, we show that the telomeric binding protein HOT1 (HMBOX1) requires dimerization – depending both on interaction with the dimerization-mediators DYNLL1/2 and its intrinsic HNFp1 domain – for its stable association with telomeres *in vivo*. Loss of HOT1 leads to an epigenetic switch from H3K9acK14ac to H3K9me3 at telomeres accompanied by attenuated ALT activity as illustrated by reduced ALT recombination by-products (C-circles) and a decrease in telomere dysfunction-induced foci (TIFs). We further show via binding assays and cross-linking mass spectrometry (XL-MS) analyses that HOT1 directly interacts with telomeric nucleosomes, providing a molecular basis for its involvement in telomeric chromatin organization. Collectively, these findings suggest that HOT1 functions as a homodimer to promote a telomeric chromatin state that is highly permissive for ALT recombination.

## Introduction

Telomeres are nucleoprotein structures that cap the ends of eukaryotic chromosomes and consist of double-stranded tandem TTAGGG repeat arrays culminating in a 3’-single-stranded overhang. Telomeres are only functional in association with the shelterin complex, comprised of the direct DNA binders TRF1, TRF2 and POT1 and the accessory factors TIN2, TPP1 and RAP1^1^. Through their unique architecture, telomeres mitigate two intrinsic challenges of linear chromosomes: (i) the need to distinguish natural chromosome ends from DNA breaks^2^, and (ii) the loss of genetic material due to the end-replication problem^3^. In somatic cells, where telomere length is not actively maintained, the end-replication problem leads to telomere attrition over successive divisions and ultimately triggers replicative senescence. In contrast, cancer cells overcome this proliferation barrier by activating telomere length maintenance mechanisms, most commonly via telomerase reactivation^4^, or in approximately 10-15% of cases through a homology-directed repair mechanism called the Alternative Lengthening of Telomeres (ALT) pathway^5–7^.

ALT-positive cells maintain telomere length by employing the break-induced replication (BIR) machinery, which orchestrates the invasion of a damaged telomere into a homologous telomeric template and its elongation via conservative DNA synthesis^8^. This telomeric elongation occurs within specialized nuclear compartments called ALT-associated PML bodies (APBs)^9–11^, which concentrate chromatin remodellers^12,13^, DNA repair factors^10,14^, and telomeric DNA^5,15^. Besides APBs, ALT-positive cells display several other defining features, including (i) long and highly heterogeneous telomeres^15^, (ii) elevated levels of telomeric sister chromatid exchanges (t-SCE)^16^, (iii) increased telomere dysfunction induced foci (TIF)^17^, (iv) elevated expression of the telomeric long noncoding RNA (TERRA)^18,19^ and (v) the accumulation of extrachromosomal telomeric repeats (e.g. C-circles, T-circles)^20,21^. Molecularly, loss-of-function mutations in the H3.3 histone chaperones ATRX and/or DAXX are among the most recurrent genetic alterations in ALT-positive cancers^22^. As a consequence, telomeres in ALT-positive cells exhibit a distinct chromatin state characterized by decreased levels of trimethylated histone H3 lysine 9 (H3K9me3) and reduced incorporation of the histone variant H3.3, contributing to a more accessible and recombination-permissive environment^19^. However, ATRX/DAXX deficiency alone does not reliably induce ALT, indicating that additional factors or changes to the telomeric chromatin are required^23–25^. Paradoxically, other studies have shown that heterochromatin may also support ALT activity. For instance, SETDB1, a histone methyltransferase that catalyses H3K9 trimethylation, has been implicated in promoting ALT by establishing a repressive chromatin environment at telomeres^13^. This has led to a model in which dynamic regulation of both chromatin decompaction and compaction may facilitate ALT, potentially at distinct stages of the pathway.

Beyond the nucleosome organization and the ubiquitous shelterin complex, telomeric chromatin is also shaped by a cohort of additional direct telomere-binders. These include ZNF524^26,27^, ZNF827^28^, NR2C2/F2^29,30^, ZBTB10^26,31^, ZBTB48^26,32,33^, and HOT1^29,34^, which collectively associate with telomeric DNA in a more dynamic and context-dependent manner. For instance, HOT1 only engages a subset of telomeres in telomerase-positive cancer cells but exhibits a markedly higher degree of telomeric association in ALT-positive cancer cells^29,32,34^, suggesting enhanced functional relevance in this context. While HOT1 was previously implicated in telomere homeostasis in both telomerase- and ALT-positive cells^34,35^, the exact molecular mechanisms remain unclear.

Here, we show that HOT1 interacts with the dimerization mediators DYNLL1 and DYNLL2. These interactions together with a HNFp1 homodimerization domain at the N-terminus of HOT1 promote HOT1 dimer formation. Disruption of either dimerization interface resulted in the loss of HOT1 binding to telomeres *in vivo*, indicating that dimerization is essential for its stable association with chromosome ends. Loss of HOT1 led to a reduction in the diacetylation mark H3K9acK14ac concurrent with increased H3K9me3 levels, reflecting a shift towards a more heterochromatic telomeric state. These chromatin changes are associated with significantly decreased C-circles and telomere dysfunction–induced foci (TIFs), indicative of reduced ALT activity. In addition, biochemical evidence shows that HOT1 directly binds telomeric nucleosomes as the underlying prerequisite for its involvement in telomeric chromatin organization. Together, these findings indicate that HOT1 contributes to the regulation of telomeric chromatin with clear implications for the regulation of ALT activity.

## Results

### Loss of HOT1 reduces ALT activity

To characterize the role of HOT1 at ALT telomeres, we first recapitulated the association of HOT1 with telomeres and APBs by immunoFISH using the PML body marker protein SP100 across five independently selected WT clones in the ALT-positive cell line U2OS. Consistent with previous observations^29,32,35^, HOT1 associated with a subset of telomeres with on average 40% of all telomeres and more than 80% of all APBs per cell (**Fig. 1A-D**). To further validate the *in vivo* association of HOT1 with ALT-positive telomeres, we performed HOT1 ChIP coupled to label-free quantitative mass spectrometry (qChIP-MS)^36^. Consistently, in qChIP-MS reactions HOT1 significantly enriched telomeric proteins and PML body-associated factors, validating that HOT1 abundantly associates with telomeres in ALT-positive cells (**Supplementary Fig. 1A, Supplementary Table 1**). Next, to test whether the high degree of association with APBs as sites of active telomere elongation represents a functional involvement of HOT1 in ALT regulation, we systematically tested the impact of HOT1 loss across ALT hallmarks. First, to assess an impact on the number of APBs, we repeated our immunoFISH analysis in five independent U2OS HOT1 KO clones. As expected, HOT1 foci were absent in the KO clones, confirming specificity of our reagents and the observed HOT1 association with ALT telomeres (**Fig. 1A**). Overall, the number of APBs per cell did not significantly differ between U2OS WT and HOT1 KO clones with on average 5.9 ± 0.7 APBs in WT cells and 5.0 ± 1.9 APBs in HOT1 KO cells (**Fig. 1E, Supplementary Fig. 1B**). Next, we evaluated TERRA expression by RT-qPCR using 5 different chromosome-arm-specific primers, but none of the TERRA transcripts showed significantly different expression between U2OS WT and HOT1 KO clones (**Supplementary Fig. 1C**). Similarly, the average telomere length measured in the five HOT1 KO clones did not vary significantly as compared to the WT clones (**Supplementary Fig. 1D**). In contrast, the frequency of telomere dysfunction induced foci (TIFs; association of DNA damage markers with telomeres as indicators of active ALT processing), single-stranded telomeric C-rich DNA (ss-TeloC) and C-circle levels were significantly reduced in U2OS HOT1 KO clones, indicating substantially attenuated ALT activity (**Fig. 1F-I)**. In sum, these data suggest that HOT1 does not impact the localisation of telomeres to APBs but participates downstream of telomere recruitment in the functional regulation of ALT activity.

**Figure 1.**
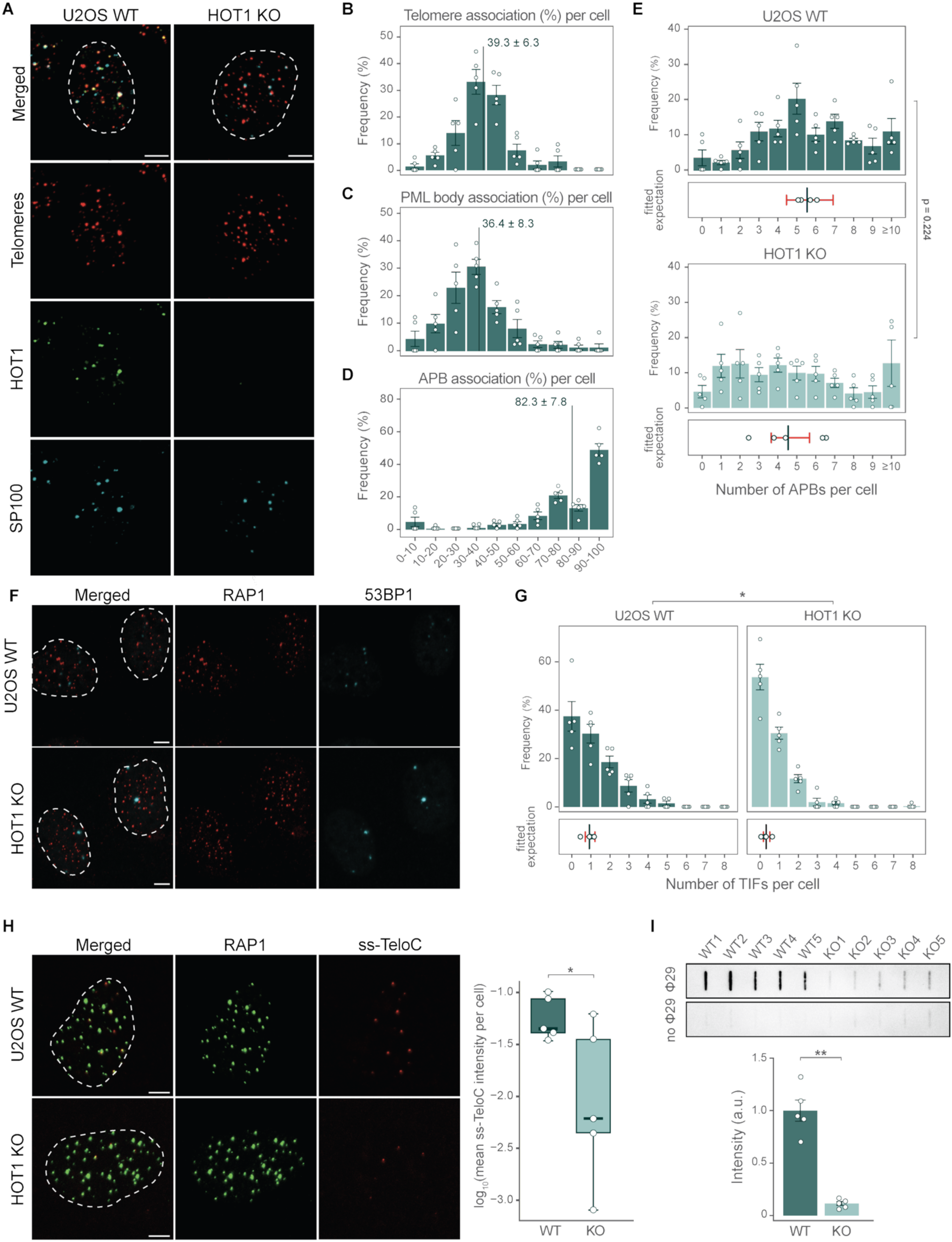
Loss of HOT1 decreases ALT activity. **(A)** Fluorescence microscopy to analyse colocalization of telomeres (red), HOT1 (green), and SP100 (cyan) in 5 U2OS WT and 5 U2OS HOT1 KO clones by immunoFISH staining. Representative images are shown (scale bar 5 µm). Dashed lines represent the nuclear outline based on counterstaining with DAPI (not shown). **(B-D)** Quantification of the frequency of colocalization events of HOT1 with **(B)** telomeres, **(C)** SP100 as a marker for PML bodies and **(D)** APBs in 5 U2OS WT clones (*n=5, 39-52 nuclei analysed per clone*). Mean values are shown with error bars representing SEM. The average % of colocalization events is indicated by black bars. **(E)** Quantification of the number of APBs per cell in U2OS WT clones (*n=5, 39-52 nuclei analysed per clone*) and U2OS HOT1 KO clones (*n= 5, 32-66 nuclei analysed per clone*). Upper plot: the frequency of cells for the indicated number of APBs, error bars represent standard error of the mean (SEM) across clones. Lower plot: the vertical line represents the fitted expected number of APBs (GLMM for negative binomially distributed data), the error bar represents the 95% confidence interval. p-value was calculated using likelihood ratio test. **(F)** Telomere-dysfunction-induced foci (TIFs) analysis in U2OS clones. Representative images of IF stainings for 53BP1 (cyan) and RAP1 (red) are shown (scale bar 5 µm). Dashed lines represent the nuclear outline based on counterstaining with DAPI (not shown). **(G)** Quantification of the number of TIFs per cell in U2OS WT clones (*n= 5, 45-54 nuclei analysed per clone*) and U2OS HOT1 KO clones (*n= 5, 46-51 nuclei analysed per clone*). Upper plot: the frequency of cells for the indicated number of TIFs, error bars represent SEM across clones. Lower plot: the vertical line represents the fitted expected number of TIFs (GLMM for negative binomially distributed data), the error bar represents the 95% confidence interval. p-value was calculated using likelihood ratio test; *p < 0.05. **(H)** ss-TeloC intensity analysis in U2OS clones. Left: representative images illustrating the immunoFISH staining of ss-TeloC (red) and RAP1 (green). Scale bar 5 µm. Dashed lines represent the nuclear outline based on counterstaining with DAPI (not shown). Right: quantification of the log-transformed ss-TeloC foci mean intensity per nuclei in U2OS WT clones (*n=5, 66-75 nuclei analysed per clone*) and U2OS HOT1 KO clones (*n=5, 63-92 nuclei analysed per clone*). Boxplots indicate the median and interquartile range; whiskers extend to 1.5x the interquartile range. p-value was calculated using Wald t-test with Satterthwaite degrees of freedom (linear mixed-effects model); *p < 0.05. **(I)** Quantification of C-circles in 5 U2OS WT and 5 HOT1 KO clones (*n=5*). C-circle reactions were carried out with 15 ng DNA template and spotted (top). Intensity for each clone was normalized by the average intensity of the 5 WT clones. Mean intensities are shown with error bars representing SD (bottom). Statistical significance was determined by Mann-Whitney test; **p < 0.01. No Φ29 indicates negative control reactions without the Φ29 DNA polymerase

### DYNLL1/2 interact with HOT1 and mediate its homodimerization

To understand how HOT1 impacts ALT activity, we first investigated HOT1 interaction partners by performing HOT1 immunoprecipitation combined with label-free quantitative mass spectrometry (IP-MS) analysis in five U2OS WT clones compared to five HOT1 KO clones as a rigorous control for antibody off-targets. In addition to enrichment of the bait protein only one protein group containing DYNLL1 and/or DYNLL2 was robustly co-immunoprecipitated with HOT1 (**Fig. 2A, Supplementary Table 2**). This interaction is not exclusive to ALT cell lines as similar experiments in HeLa cells also identified DYNLL1/2 as the primary interaction partner of HOT1 (**Supplementary Fig. 2A, Supplementary Table 2**). Due to very high sequence identity between both paralogs these initial mass spectrometry data could not distinguish between enrichment of DYNLL1, DYNLL2 or both. To differentiate between these possibilities and to further validate the putative HOT1-DYNLL1/2 interaction, we performed reciprocal IP-MS experiments with FLAG-tagged expression constructs. Both individual IPs with DYNLL1 and DYNLL2 independently enriched HOT1, suggesting that HOT1 can interact with both paralogs. In addition, we identified several known DYNLL1/2 interaction partners such as members of the dynein complex (DYNC1I2, DYNC1H1 and DYNC1L1/2)^37^ and TP53BP1^38,39^ (**Fig. 2B, Supplementary Fig. 2B-C, Supplementary Table 3**). Again, these reciprocal interactions are not limited to ALT-positive cells and were confirmed in HeLa cells^40^ (**Supplementary Fig. 2D-E, Supplementary Table 3**). The much larger number of DYNLL1/2 interaction partners is not surprising since DYNLL1/2 are small homo-dimeric proteins that have previously been reported to function as a scaffold, mediating the protein dimerization of their interaction partners^41^. DYNLL1/2 are known to interact with a short recognition sequence, which is most frequently anchored by a TQT-motif^42^. Indeed, HOT1 contains an N-terminal TQT motif that is predicted by the DYNLL1/2 binding partner prediction tool (LC8Pred tool^43^) to mediate interaction with DYNLL1/2 (**Fig. 2C**). To test this, we reintroduced either FLAG-HOT1 WT or FLAG-HOT1ΔTQT in four U2OS HOT1 KO clones and immunoprecipitated FLAG-HOT1. In agreement with the above, FLAG-HOT1 WT robustly co-immunoprecipitated both DYNLL1 and DYNLL2 compared to FLAG-HOT1ΔTQT, demonstrating that their interaction indeed requires the HOT1 TQT motif (**Fig. 2D, Supplementary Fig. 2F-G, Supplementary Table 4**).

**Figure 2.**
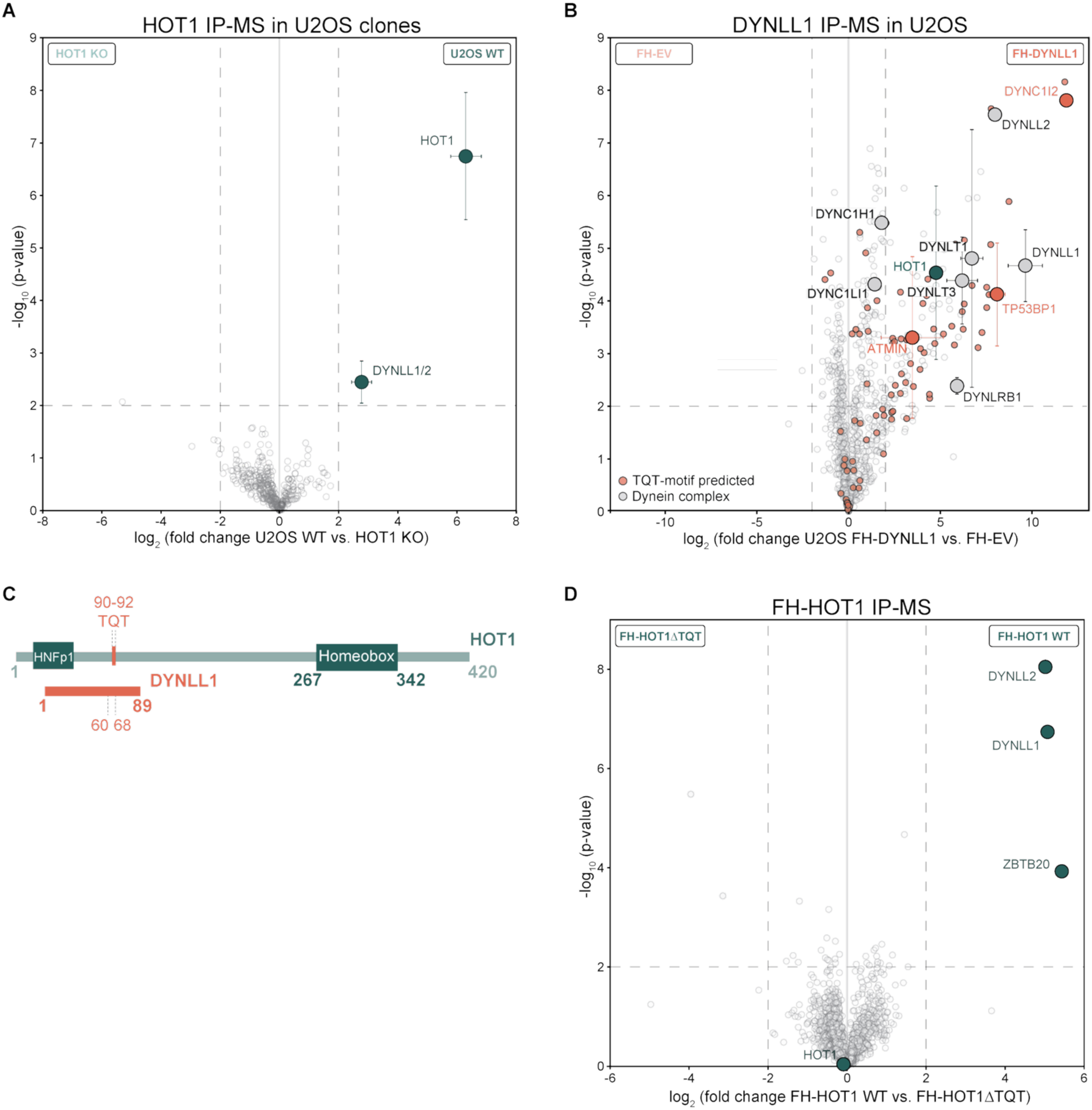
DYNLL1/2 interact with HOT1 via its TQT-motif. **(A)** Volcano plot comparing HOT1 IP-MS reactions from 5 U2OS WT clones and 5 U2OS HOT1 KO clones (*n=5*). **(B)** Volcano plot comparing FLAG IP-MS reactions from U2OS cells expressing either FLAG-DYNLL1 or FLAG empty vector (EV) (*n=4*). Proteins predicted to interact with DYNLL1 based on the LC8Pred tool^43^ are highlighted in red and members of the dynein complex are highlighted in grey. **(C)** Schematic representation of the interaction between HOT1 (green) and DYNLL1/2 (red) with HOT1’s TQT motif highlighted in red. **(D)** Volcano plot comparing FLAG IP-MS reactions from 4 U2OS HOT1 KO clones expressing either FLAG-HOT1 WT or FLAG-HOT1ΔTQT (*n=4*). **(A-B, D)** All pull-downs were analysed by label-free quantification. Specifically enriched proteins are distinguished from background binders by a two-dimensional cut-off of > 4-fold enrichment and p-value < 0.01. Two-dimensional error bars represent the standard deviation based on iterative imputation cycles during the label-free analysis to substitute missing values (e.g. no detection in the HOT1 KO control)

### HOT1 dimerization is necessary for its association with telomere *in vivo*

We had originally identified HOT1 via *in vitro* reconstitution DNA pull-down assays and we were able to generate a co-crystal structure of the HOT1 homeobox domain with telomeric DNA in the absence of DYNLL1/2^34^, suggesting that *in vitro* binding to telomeric DNA is independent of putative dimerization. However, given that the shelterin proteins TRF1 and TRF2 also only require their homeobox domain for *in vitro* binding^44^ but are dependent on dimerization for their *in vivo* telomere association^45^, we hypothesised that the same might be true for HOT1. To test whether the DYNLL1-mediated HOT1 dimerization impacts its association to telomeres, we depleted DYNLL1 in U2OS cells with shRNAs (**Supplementary Fig. 3A**) and assessed co-localisation of HOT1 with TRF2 as a telomeric marker by immunofluorescence staining. Overall, we observed a reduction in the number of HOT1 foci upon DYNLL1 knock-down with both shRNAs (**Fig. 3A-B**). In addition, we observed a concomitant decrease in HOT1 association with telomeres with on average 21.5% and 15.9% of telomeric foci associated with HOT1 for DYNLL1 shRNAs #1 and #2, respectively, compared to 40.2% colocalization with shGFP control (**Fig. 3C**). Consistent with the above, re-expression of HOT1ΔTQT constructs in HOT1 KO cells resulted in a marked decrease in cells with HOT1 foci despite comparable expression to the WT construct (**Fig. 3D-E, Supplementary Fig. 3B**). In contrast, HOT1 WT rescue restored foci formation in a substantial fraction of cells with a foci distribution profile resembling endogenous HOT1 (**Fig. 3D-E, Supplementary Fig. 3B**).

**Figure 3.**
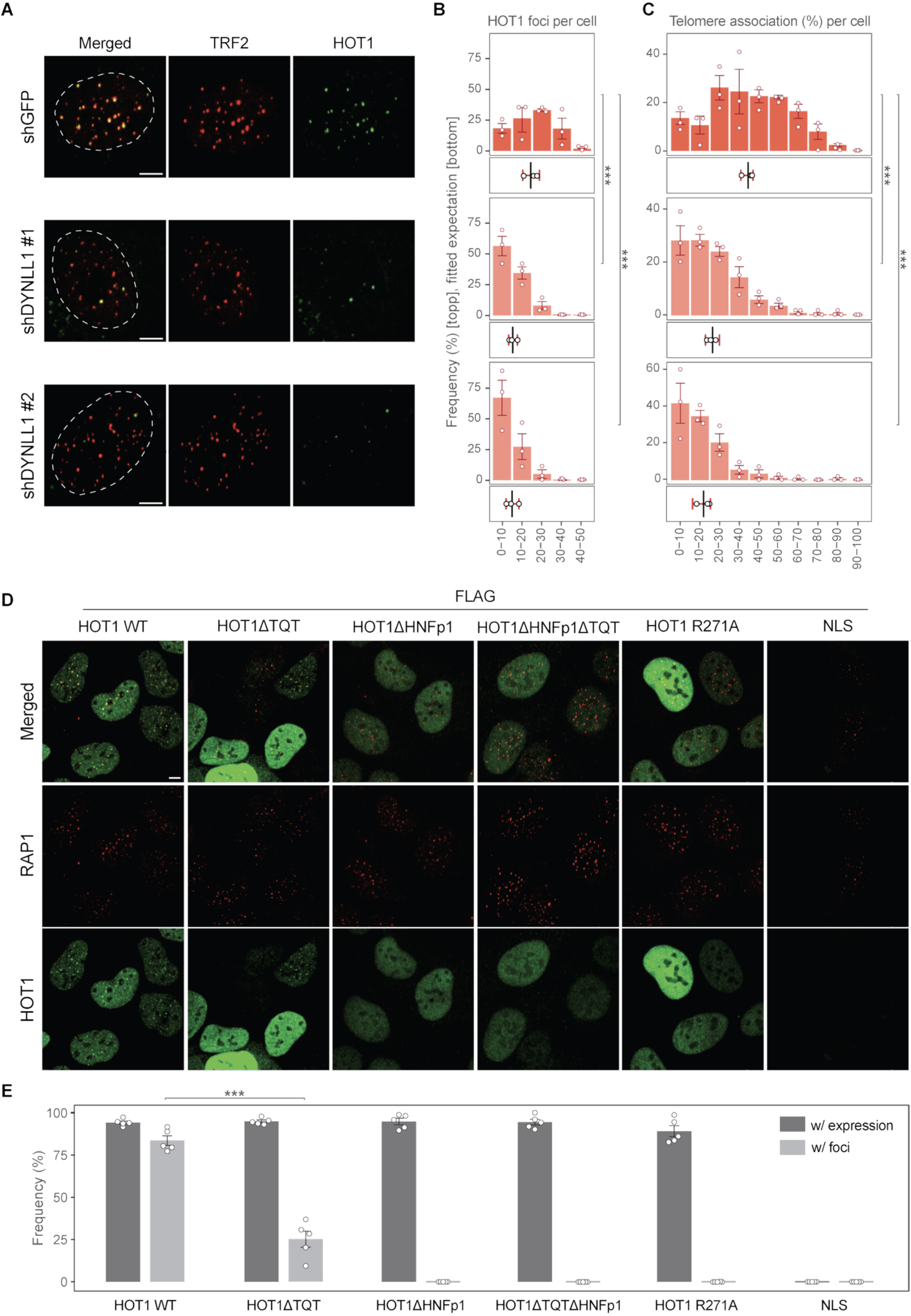
HOT1 dimerization is necessary for telomere binding *in vivo*. **(A)** Fluorescence microscopy to analyse colocalization between HOT1 (green) and TRF2/telomeres (red) in U2OS cells with shGFP vs. shDYNLL1 #1 and #2 by immunofluorescence staining (*n=3*). Representative images shown (scale bar 5 µm). Dashed lines represent the nuclear outline based on counterstaining with DAPI (not shown). **(B-C)** Quantification of **(B)** the number of HOT1 foci per cell and **(C)** the percentage of HOT1-positive telomeres in U2OS shGFP (*n=3, 71-91 nuclei per replicate*), U2OS shDYNLL1 #1 (*n=3, 72-94 nuclei per replicate*) and U2OS shDYNLL1 #2 (*n=3, 63-102 nuclei per replicate*). Upper plot: the frequency of cells for the indicated number of **(B)** HOT1 foci per cell or **(C)** the percentage of telomeres association per cell, error bars represent SEM across clones. Lower plot: the vertical line represents the fitted expected number of HOT1 foci and percentage of telomere associations (CLMM for ordinally distributed data), the error bar represents the 95% confidence interval. p-values were calculated using Wald test of the CLMM coefficient; ***p < 0.001. **(D)** Colocalization analysis as in (**A**) in 5 U2OS HOT1 KO clones expressing either FLAG-HOT1 WT, FLAG-HOT1ΔTQT, FLAG-HOT1ΔHNFp1, FLAG-HOT1ΔHNFp1ΔTQT, FLAG-HOT1 R271A and FLAG-NLS. **(E)** Quantification of the percentage of cells expressing the HOT1 constructs and the percentage of cells with foci in U2OS FLAG-HOT1 WT (*n=5, 79-110 nuclei per replicate*), U2OS FLAG-HOT1ΔTQT (*n=5, 78-106 nuclei per replicate*), U2OS FLAG-HOT1ΔHNFp1 (*n=5, 72-96 nuclei per replicate*), U2OS FLAG-HOT1ΔHNFp1ΔTQT (*n=5, 39-75 nuclei per replicate*), U2OS FLAG-HOT1 R271A (*n=5, 43-97 nuclei per replicate*) and U2OS FLAG-NLS (*n=5, 56-87 nuclei per replicate*). Error bars represent standard error of the mean (SEM) across clones (*n=5*) and p-value was calculated using Wald z-test (GLMM for negative binomially distributed data); ***p < 0.001. Each clone is indicated by a dot.

Interestingly, HOT1 is predicted to contain an HNFp1 domain, which in other members of the same protein family, HNF1α and HNF1β, is known to mediate homodimerization^46^. To explore whether the HNFp1 domain may additionally contribute to telomeric localization, we reintroduced HOT1ΔHNFp1 and HOT1ΔHNFp1ΔTQT mutants into HOT1 KO cells. Both mutants showed a complete loss of HOT1 foci, comparable to the DNA binding-deficient R271A mutant^34^ (**Fig. 3D**). Taken together, these findings support a model in which proficient binding of HOT1 to telomeres *in vivo* depends on both DYNLL1/2- and HNFp1-mediated dimerization.

### Loss of HOT1 impacts the telomeric chromatin composition

While the above indicates an intricate regulation of *in vivo* binding to telomeric chromatin, the HOT1-DYNLL1/2 interaction *per se* does not explain how HOT1 affects ALT activity. Therefore, we next assessed changes in the telomeric chromatin composition in the presence or absence of HOT1 by chromatin immunoprecipitation coupled to label-free quantitative mass spectrometry (qChIP-MS)^36^ using TRF2 as bait protein. When comparing TRF2 qChIP-MS reactions of each five U2OS WT and HOT1 KO clones we identified moderate differential enrichment of the chromatin remodeller CHD3 at telomeric chromatin of WT cells and ORC1 at telomeric chromatin of HOT1 KO cells, respectively (**Fig. 4A, Supplementary Fig. 4A-B, Supplementary Table 5**). In parallel, we explored potential changes in epigenetic marks in the absence of HOT1 and interrogated our mass spectrometry data for mono-, di-, tri-methylation and acetylation on histone peptides. While we were not able to detect histone methylation sites in our analysis, we identified di-acetylated H3K9acK14ac peptides with a robust >10-fold reduction in U2OS HOT1 KO clones compared to U2OS WT clones (**Fig. 4B, Supplementary Table 5**). We validated these results by performing reciprocal ChIP slot blot reactions in 5 U2OS WT and HOT1 KO clones. Consistent with the mass spectrometry results, more telomeric DNA was enriched in U2OS WT as compared to U2OS HOT1 KO clones using an anti H3K9acK14ac antibody while global histone H3 and mono-H3K9ac levels were unaffected (**Fig. 4C-D**). Of note, the mutually exclusive H3K9me3 modification enriched more telomeric DNA in the U2OS HOT1 KO clones, suggesting a molecular switch from the activating H3K9acK14ac mark in U2OS WT cells to the repressive H3K9me3 mark in U2OS HOT1 KO clones (**Fig. 4C-D**). The above results support a model in which HOT1-dependent ALT activity regulation is driven by changes in telomeric chromatin accessibility upstream of telomeric recombination.

**Figure 4.**
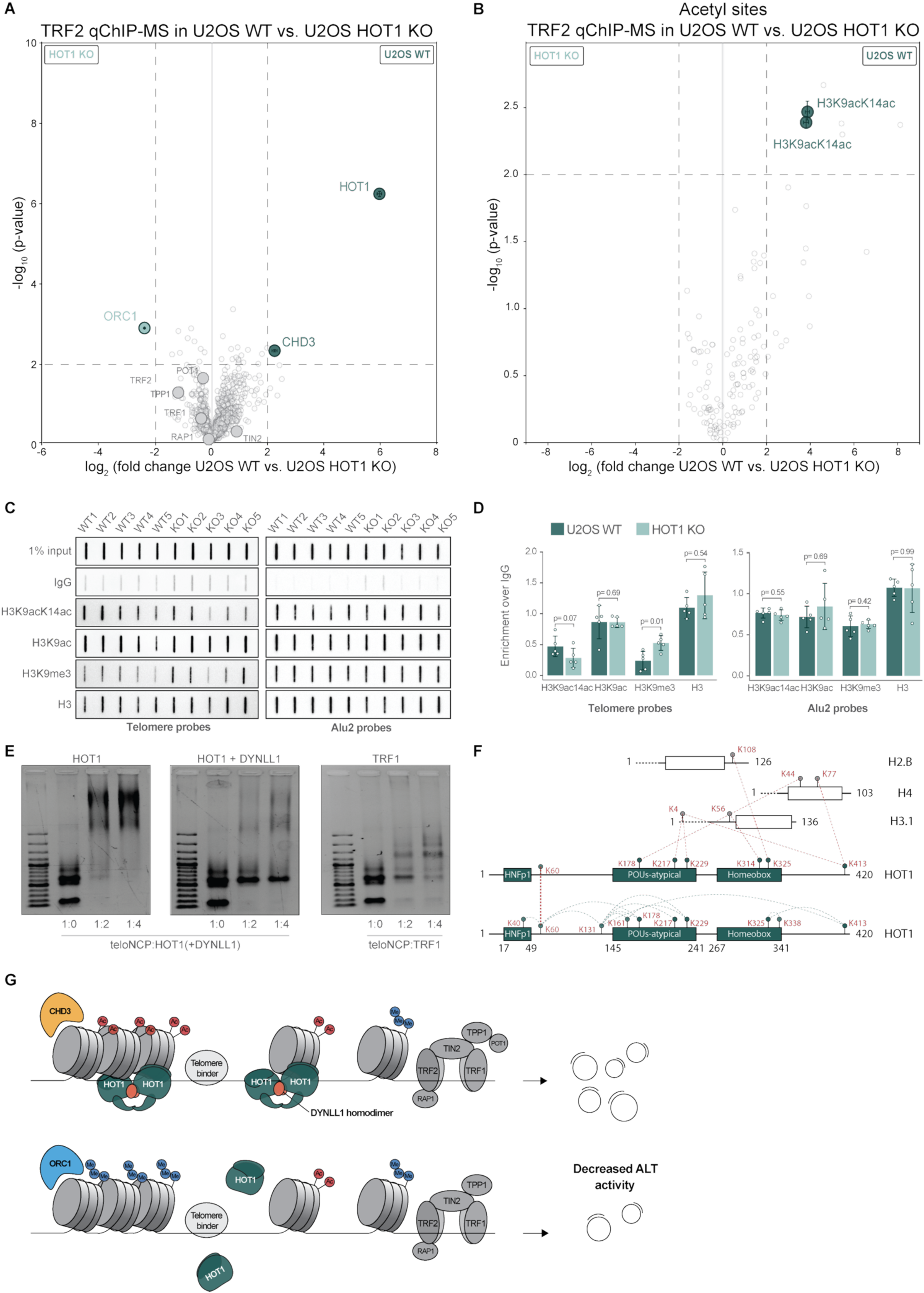
Loss of HOT1 impacts telomeric chromatin composition. **(A-B)** Volcano plots comparing TRF2 qChIP-MS reactions analysed by label-free quantitation in 5 U2OS WT clones vs. 5 U2OS HOT1 KO clones (*n=5*). Enriched proteins **(A)** and enriched acetyl (K) sites **(B)** are distinguished from background by a two-dimensional cut-off with fold enrichment > 4 and p-value < 0.01. Two-dimensional error bars represent the standard deviation based on iterative imputation cycles during the label-free analysis to substitute missing values (e.g. no detection in the HOT1 KO control). Proteins enriched at telomeres in U2OS WT and HOT1 KO clones are indicated in dark green and light green, respectively. **(C)** Histone H3 post-translational modifications and IgG ChIP assays were performed in 5 U2OS WT and HOT1 KO clones (*n=5*). Telomeric or Alu2 repeats were detected by dot blotting and hybridization with (CCCTAA)4 telomeric (left) or Alu2 probes (right). **(D)** Quantification of ChIP slot blot reactions for the various H3 post-translation modifications. Mean intensities normalized against IgG are shown with error bars representing SD. Statistical significance was determined by Mann-Whitney test and p-values are indicated on the graphs. **(E)** Electrophoretic mobility assay of telomeric mononucleosomes bound with HOT1 (left) or HOT1+DYNLL1 (middle) at the indicated NCP:protein molar ratios. Corresponding data with TRF1 (right) serve as a positive control. **(F)** Representative map of crosslinks detected by XL-MS analysis between HOT1 and telomeric NCP. Interactions between HOT1 and histones are displayed in red dashed lines. Interactions within HOT1 or between two HOT1 monomers are highlighted in green dashed lines. **(G)** Schematic summary of our study.

These results further raise the question whether HOT1 may be able to engage telomeric DNA in the context of chromatin. To address this, we carried out electrophoretic mobility shift assays (EMSA) to investigate HOT1 and HOT1-DYNLL1 interaction with reconstituted telomeric mono- and di-nucleosomes. Indeed, both purified HOT1 alone and in association with DYNLL1 interacted with the telomeric mono- and di-nucleosome similarly to TRF1, suggesting that HOT1 can directly bind to telomeric chromatin (**Fig. 4E and Supplementary Fig. 4C**). Interestingly, HOT1 alone showed a distinct higher order band in comparison to a diffused banding pattern in presence of HOT1-DYNLL1 in complex with telomeric nucleosome core particles (NCP). To examine this and validate that there is no histone eviction upon HOT1 binding to telomeric nucleosomes, we carried out MgCl2 mediated precipitation of HOT1 and HOT1-DYNLL1 complexed with telomeric nucleosomes. In the presence of MgCl2, the factors that are bound to nucleosomes precipitate along with histones due to charge neutralization by MgCl2 and unbound factors remain in the supernatant fraction. Indeed, HOT1 co-precipitated with nucleosomes, supporting HOT1 association with telomeric nucleosomes (**Supplementary Fig. 4D**). Consistent with the EMSA results, cross-linking mass spectrometry (XL-MS) analysis of HOT1-NCP complexes identified multiple cross links between HOT1 and H2B, H3 and H4 histone proteins. Interestingly, in addition to crosslinks in the homeobox domain, we identified recurrent contacts involving residues in the adjacent predicted POUs-atypical domain (**Fig. 4F, Supplementary Table 6**). In parallel, our XL-MS dataset highlighted an extensive network of inter- and intramolecular cross-links within HOT1 mostly concentrated around the predicted HNFp1 and the POUs-atypical domains. Of note, we identified a symmetric cross-link between two equivalent K60 residues (**Fig. 4F, Supplementary Table 6**), confirming that HOT1 homodimerizes via its N-terminal region. Having established that both HOT1 and HOT1-DYNLL1 engage telomeric chromatin, we performed cryo-EM screens to further analyse the HOT1-NCP engagement and identified clearly visible features based on isolated nucleosomes with additional density as well as aggregates for both NCP-HOT1 and NCP-HOT1-DYNLL1 complexes (**Supplementary Fig. 4E-F**). The presence of aggregates indicates the propensity of the complex to oligomerize. It is noteworthy that the HOT1-NCP complex showed comparatively less aggregation (**Supplementary Fig. 4F**), suggesting that the NCP-HOT1-DYNLL1 complex may yield more pronounced oligomerization, visualized as clutches of nucleosomes.

Collectively, our results support a model in which HOT1 contributes to ALT activity by influencing the telomeric chromatin organization (**Fig. 4G**). In such a model, HOT1 requires dimerization to directly engage telomeric chromatin and to prevent the accumulation of H3K9me3 at telomeres.

## Discussion

HOT1 is a telomere-associated protein previously identified as a positive telomere length regulator in telomerase-positive cells^34,47^, yet accumulating evidence points to a more prominent role in the ALT context. HOT1 displays higher abundance at ALT telomeres compared to telomerase-positive cells^29,32,35^, and it localizes to nearly all ALT-associated PML bodies (APBs), positioning it as part of the sites of recombination-driven telomere elongation. Here, our findings reveal that HOT1 influences the telomeric chromatin environment and supports ALT activity. Loss of HOT1 resulted in a marked reduction in C-circles without affecting APB formation, suggesting that HOT1 acts downstream of APB assembly during ALT recombination. This view is further supported by concomitant reduction in ss-TeloC signals and a decrease in the number of telomere dysfunction-induced foci (TIFs), both consistent with diminished ALT activity.

In addition to reduced ALT activity, HOT1-deficient cells displayed reduced H3K9acK14ac and elevated H3K9me3 levels at telomeres while H3K9ac levels remained similar to WT cells. ALT telomeres are thought to exist in a heterochromatin-leaning state, with substantial presence of H3K9me3 yet sufficient local accessibility to sustain recombination-based elongation. In this context, the increase in telomeric H3K9me3 and the concomitant reduction in H3K9acK14ac that we observe upon HOT1 loss point to a switch towards a more stably repressed conformation. Because the H3K9ac-specific antibody recognizes H3K9ac only when K14 is unmodified, the stable H3K9ac signal together with the reduced H3K9acK14ac signal indicates a selective depletion of the di-acetylated histone population rather than a global loss of H3K9ac. This pattern is consistent with the idea that, in the absence of HOT1, H3K9ac in the context of H3K14ac preferentially transitions towards H3K9me3, although our data do not determine whether H3K14ac is maintained or removed during this process. Interestingly, the H3K9acK14ac mark has been linked to bivalent promoters, where it defines a poised state that can be dynamically shifted towards activation or repression depending on the cellular context. In particular, one study reported that H3K9me3/H3K14ac dually marked regions define a poised inactive state that can be promptly activated upon the loss of one or both chromatin marks^48^. If a similar poised configuration exists at ALT telomeres, the switch upon HOT1 loss may compromise the rapid, recurrent access of DNA-damage and recombination factors required to sustain efficient ALT activity.

It is particularly intriguing how HOT1 influences telomeric chromatin organization to support ALT activity. One potential insight comes from HOT1’s ability to homodimerize. HOT1 contains a predicted HNFp1 homodimerization domain as well as a TQT-motif which mediates its interaction with DYNLL1/2. The coexistence of intrinsic and DYNLL1/2-mediated dimerization suggests a bipartite homodimerization process, whereby DYNLL1/2 stabilizes HOT1 self-assembly and may promote higher-order oligomer formation. Our data show that the deletion of either DYNLL1-mediated dimerization or the predicted intrinsic dimerization interface significantly reduced HOT1 foci formation at telomeres, highlighting the importance of HOT1 homodimerization for its stable association with telomeric chromatin. This behaviour is reminiscent of TRF1 and TRF2, whose TRFH-mediated dimerization is dispensable for sequence-specific DNA binding but essential for stable telomeric assembly and function^45^. Building on this, our biochemical assays further indicate that HOT1 can directly engage telomeric chromatin: HOT1 binds reconstituted telomeric nucleosomes in EMSA, and XL-MS identifies multiple cross-links between HOT1 and the histone core, with recurrent contacts mapping to the predicted POUs-atypical region as well as additional sites in the homeobox and C-terminal regions. It is tempting to speculate that the HOT1-nucleosome engagement may structurally mask access for readers and/or writers and thus prevents conversion from H3K9acK14ac to H3K9me3. High resolution cryo-EM structures of HOT1 with telomeric nucleosomes and DYNLL1/2 should address this in the future.

Finally, HOT1-deficient cells showed reduced CHD3 and increased ORC1 enrichment at telomeres. CHD3, an ATP-dependent chromatin remodeller within the NuRD complex, is broadly linked to nucleosome repositioning and deacetylation^49^. In contrast, ORC1 can bind to H3K9me3 and promote H3K9me3-associated chromatin repression^50^, and also has been linked to telomeric H3K9 methylation^51^. The opposing changes in CHD3 and ORC1 abundance therefore further support that HOT1 acts upstream of a chromatin remodelling switch. Whether these changes arise directly from the loss of HOT1 or secondarily from altered telomeric chromatin architecture remains to be determined, but they provide a mechanistic framework, linking HOT1 depletion to reduced recombination permissiveness.

Taken together, our findings suggest a model in which HOT1 participates in shaping the epigenetic landscape at ALT telomeres. We hypothesize that, by forming higher-order assemblies through its bipartite dimerization mechanism engaged with telomeric nucleosomes, HOT1 helps maintain a chromatin environment that is inherently accessible for recombination-based telomere elongation. Loss of HOT1 shifts telomeric chromatin towards a more heterochromatic, less recombination-permissive state, resulting in reduced ALT activity.

## Materials and Methods

### Cell culture

U2OS (CVCL_0042) cells were cultivated in Dulbecco’s Modified Eagle Medium (DMEM) containing 4.5 g/L glucose, 4 mM glutamine, 1 mM sodium pyruvate and supplemented with 10% fetal bovine serum (Gibco), 100 U/mL penicillin and 100 µg/mL streptomycin (Gibco). Cells were kept at 37 °C and 5% CO2 in a humidified incubator. Cell lines were authenticated by STR profiling (1st BASE) and routinely monitored for mycoplasma contamination using the MycoAlert PLUS mycoplasma detection kit (Lonza).

### Cloning and plasmids

Full-length HOT1 was PCR amplified from a previously established pcDNA3.1 vector^34^ while DYNLL1 and DYNLL2 were amplified from U2OS cDNA (see quantitative real-time PCR section) using the Q5 Hot Start DNA polymerase (NEB) according to manufacturer’s conditions. HOT1, DYNLL1 and DYNLL2 inserts were cloned using XhoI and NotI restriction digestion into a modified pOZ-FH-N vector with a puromycin or neomycin resistance cassette inserted in lieu of the IL-2 receptor selectable marker by PCR swap. HOT1 mutations were introduced by Q5 site-directed mutagenesis (NEB) according to manufacturer’s conditions. For gene knock-down, short hairpin RNAs (shRNAs) were shortlisted from the Broad Institute GPP web portal (https://portals.broadinstitute.org/gpp/public/gene/search) and were cloned into the pLKO.1-puro vector. Briefly, 5 µg of pLKO.1 were linearized using AgeI/EcoRI (NEB) restriction enzyme digestion. Complementary shRNA oligonucleotides (100µM) were first annealed in NEB buffer 2 (4 min at 95°C followed by 10 min at 70°C, followed by gradual cooling to room temperature for approximately 2 h), phosphorylated using 10 U of T4 PNK (NEB) at 37°C for 30 min then purified with Qiagen MinElute PCR purification kit according to manufacturer’s conditions. 25 ng of digested vector and 5 ng of purified shRNA oligonucleotides were then ligated overnight at 16°C using 10 U T4 ligase (Thermo). All primers used for cloning are listed in **Supplementary Table 7**. Plasmid sequences were confirmed by Sanger sequencing (1st BASE) and/or nanopore whole plasmid sequencing (Plasmidsaurus).

### Transfection and transduction

For retrovirus production, 300,000 HEK293T cells were transfected using 1 µg of the retroviral transfer vector, 0.25 µg of the packaging vector pMD-MLV and 0.25 µg of the envelope vector pCMV-VSV-G using 2.5 µg polyethylenimine (PEI, MW 25,000; Polysciences) diluted in Opti-MEM (Gibco). For lentivirus production, 0.5 µg of the transfer vector was co-transfected with 0.25 µg each of the packaging vectors pRSV-Rev and pMDLg/pRRE, along with 0.25 µg of the envelope vector pMD2.G using 2.5 µg polyethylenimine (PEI, MW 25,000; Polysciences) diluted in Opti-MEM (Gibco). Viral supernatants were collected and filtered through a 0.45 µm membrane 3 days post-transfection. For transduction, 150,000 cells were inoculated for 24 h with 2 mL viral supernatants supplemented with 10 mM HEPES and 8 µg/mL polybrene (Sigma). 48 h post-inoculation, cells were selected using 0.8 µg/mL puromycin (Gibco) or 400 µg/mL neomycin (Gibco) for 3 and 5 days, respectively.

### Quantitative real-time PCR

Total RNA was extracted using the RNeasy Plus Mini Kit (Qiagen) according to the manufacturer’s instructions. For TERRA expression analysis, the optional on-column DNase I treatment (Qiagen) was included. For cDNA synthesis, 1 μg of total RNA was reverse transcribed using the SuperScript IV First-Strand Synthesis System (Invitrogen) with oligo(dT) or TERRA-specific primers (**Supplementary Table 7**) according to the manufacturer’s guidelines. Quantitative PCR was performed using 1x QuantiNova SYBR Green PCR Master Mix (Qiagen) with 500 nM of each primer (**Supplementary Table 7**) on a QuantStudio 5 Real-Time PCR system (Thermo Scientific). Thermal cycling conditions were as follows: initial denaturation at 95 °C for 2 min, followed by 40 cycles of 95 °C for 5 sec and 60 °C for 30 sec, ending with a melting curve analysis. Ct (cycle threshold) values of individual target genes were normalized to TBP, which served as the housekeeping gene.

### Southern blot analysis of Telomere Restriction Fragment (TRF) length

U2OS genomic DNA was extracted using the QIAamp DNA Blood Mini Kit (Qiagen) according to the manufacturer’s instructions. TRF length analysis was performed using the TeloTTAGG telomere length assay kit (Roche) according to manufacturer’s instructions with slight modifications. Briefly, 7-10 µg of genomic DNA was digested overnight using 20 U of HinfI and RsaI (NEB) at 37 °C. Digested genomic DNA was then resolved for 17 h on a 1% low-melt agarose gel (Bio-Rad) with an initial switch time of 0.1 s and a final switch time of 3.36 s at 6 V/cm (Bio-Rad) in 1x TAE buffer. Following electrophoresis, the gel was incubated in 0.25 M HCl for 30 min for depurination, then rinsed twice with distilled water. Subsequently, the gel was incubated twice in denaturation solution (0.5 M NaOH, 1.5 M NaCl) for 20 min each, rinsed twice with distilled water followed by incubation in neutralizing solution (0.5 M Tris-HCl, 3 M NaCl, pH 7.5) twice for 20 min each. The DNA was then transferred overnight to a positively charged nylon membrane (Hybond, N+, Amersham) by capillary osmosis in presence of 20x SSC (3 M NaCl, 0.3 M sodium citrate tribasic dehydrate, pH 7) and fixed by UV-crosslinking at 120 mJ using a Stratalinker UV Crosslinker (Stratagene). The membrane was then rinsed twice with distilled water and incubated in pre-warmed DIG Easy Hyb Granules provided in the kit for one hour at 42 °C. The membrane was then hybridized with DIG-labelled telomere probe (1:5,000) for 3 h at 42 °C. The membrane was washed twice with stringent buffer 1 (2x SSC, 0.1% SDS) for 5 min each, twice with pre-warmed stringent buffer 2 (0.2x SSC, 0.1% SDS) at 50 °C for 15 min each and rinsed with 1x wash buffer provided in the kit for 5 min. The membrane was blocked with 1x blocking solution for 30 min, followed by incubation with anti-DIG-AP antibody (1:10,000) diluted in blocking solution for 30 min. Subsequently, the membrane was washed twice with 1x wash buffer for 15 min and incubated in 1x detection solution for 5 min. The telomeric DNA was detected using the digoxigenin luminescent detection (CDP star) system for 5 min at room temperature and imaged on a ChemiDoc MP Imaging System (Bio-Rad). Average telomere length was calculated by comparison to the DNA ladder provided in the kit using WALTER (Web-based Analyser of the Length of Telomeres)^52^.

### C-circle assay

Genomic DNA was extracted using the QIAamp DNA Blood Mini Kit (Qiagen) according to the manufacturer’s instructions. 300 ng of genomic DNA was digested using 10 U of HinfI and RsaI (NEB) each at 37 °C for 2 h. Subsequently, 15 ng of digested DNA was amplified with 7.5 U of Φ29 polymerase (NEB) in 1x Φ29 buffer supplemented with 0.1 mg/mL recombinant BSA and 2 mM dATP, dTTP, dGTP for 6 h at 30 °C. After heat inactivation (20 min at 70°C), amplified products were diluted in 2x SSC and slot-blotted on a positively charged nylon membrane (Hybond H+, Amersham) and UV cross-linked at 120 mJ using a Stratalinker UV Crosslinker (Stratagene). Reactions lacking Φ29 polymerase served as negative controls. C-circle assay analysis was performed using the TeloTTAGG telomere length assay kit (Roche) as described in the “Southern blot analysis of Telomere Restriction Fragment (TRF) length” section.

### Immunofluorescence and fluorescence in situ hybridization (immunoFISH) staining

Cells were seeded 24 h prior to the immunofluorescence staining procedure on glass coverslips (VWR). Cells were washed once with 1x PBS, fixed in 10% formalin solution (Sigma-Aldrich) for 10 min and rinsed with 1x PBS complemented with 30 mM glycine (30 mM glycine-PBS). Permeabilization was performed using 1x PBS containing 0.5% Triton X-100 for 5 min at 4 °C, followed again by two washes with 30 mM glycine-PBS. Cells were then blocked for 30 min in blocking buffer (0.2% fish skin gelatin (Sigma-Aldrich) in 1x PBS). Primary antibodies were incubated for 1 h (**Supplementary Table 8**) then washed 3 times for 3 min in blocking buffer. Next, Alexa Fluor coupled secondary antibodies (**Supplementary Table 8**) were incubated for 30 min in the dark and then washed once for 3 min in blocking buffer. To stain nuclei, 1 µg/mL DAPI solution (Thermo Scientific) was applied for 3 min, then washed three times for 3 min each in blocking buffer. Coverslips were mounted using ProLong Gold Antifade Reagent (Thermo Scientific).

When telomeric FISH was combined with immunofluorescence staining, FISH staining was carried out immediately after the secondary antibody incubation step in the above protocol. Cells were post-fixed for 10 min with 10% formalin solution, then rinsed three times with 30 mM glycine-PBS. Next, cells were dehydrated consecutively with 70%, 90% and 100% EtOH washes for 5 min each and coverslips were air-dried for at least 10 min. 1x telomeric PNA probes (Panagene) were denatured for 5 min at 70 °C, centrifuged at maximum speed and then diluted to 1x in 70% formamide solution and dispensed to each coverslip. Coverslips were heated up to 80 °C for 10 min to denature DNA and probes were allowed to hybridize for 90 min in the dark. After hybridization, coverslips were washed twice for 15 min each with 70% formamide solution and four times for 5 min with Tris/NaCl/Tween-20 solution (50 mM Tris pH 8, 150 mM NaCl and 0.05% Tween-20). Subsequently, cells were once again dehydrated consecutively with 70%, 90% and 100% EtOH washes for 2 min each and coverslips were air-dried. DAPI staining and mounting were performed as indicated above.

Images were acquired with a Leica SP8 or a Zeiss LSM 880 AiryScan confocal microscope using a 100x oil immersion objective and were processed using Zen Pro software (Zeiss). Z-stacks were deconvolved and subsequently reconstructed as z-projections using FIJI (Image J).

### ss-TeloC assay

The ss-TeloC assay was combined with immunofluorescence staining similar to immunoFISH stainings and was carried out immediately after the secondary antibody incubation step. Cells were seeded on cell culture chamber slides (ibidi) 24 h prior to the immunofluorescence staining procedure. Following telomeric immunofluorescence staining, cells were post-fixed 10 min at room temperature with 10% formalin solution, then rinsed three times with 30 mM glycine-PBS. Next, cells were dehydrated consecutively with 70%, 90% and 100% EtOH washes for 5 min each and coverslips were air-dried for at least 10 min. 1x G-rich telomeric PNA probes (Panagene) were denatured for 5 min at 70 °C, centrifuged at maximum speed and then diluted to 1x in 50% formamide solution and dispensed to each coverslip. Probes were allowed to hybridize for 90 min in the dark. After hybridization, coverslips were washed four times for 5 min with Tris/NaCl/Tween-20 solution (50 mM Tris pH 8, 150 mM NaCl and 0.05% Tween-20). Subsequently, cells were once again dehydrated consecutively with 70%, 90% and 100% EtOH washes for 2 min each and coverslips were air-dried. DAPI staining and mounting were performed as in the immunofluorescence staining section.

### Nuclear and whole cell protein extraction

To prepare whole cell protein extracts, cells were pelleted and washed with 1x PBS and then incubated in RIPA lysis buffer (Sigma-Aldrich) supplemented with 1x cOmplete protease inhibitor cocktail (Roche) for 30 min on ice. Whole cell protein lysate was collected following a 15 min centrifugation at 20,000 g. To prepare nuclear extracts, cells were collected as described above and then incubated for 10 min on ice in a hypotonic buffer (10 mM HEPES (pH 7.9), 1.5 mM MgCl₂, 10 mM KCl and 1x cOmplete protease inhibitor cocktail (Roche)). Cell lysis was performed mechanically (30-45 strokes) using a dounce homogenizer containing hypotonic buffer supplemented with 0.1% IGEPAL CA-630 (Sigma-Aldrich) and 0.5 mM DTT (Sigma-Aldrich). Isolated nuclei were washed once with 1x PBS and subsequently extracted in a hypertonic buffer (420 mM NaCl, 20 mM HEPES pH 7.9, 20% (v/v) glycerol, 2 mM MgCl₂, 0.2 mM EDTA, 0.1% IGEPAL CA-630, 0.5 mM DTT and 1x cOmplete protease inhibitor cocktail (Roche)) for 2 h at 4 °C on a rotating wheel. Nuclear extracts were collected following 1 h centrifugation at maximum speed. Protein quantification was carried out using the Pierce BCA Protein Assay kit (Thermo Scientific) according to the manufacturer’s instructions.

### Protein immunoprecipitation

Dynabeads Protein G magnetic beads (Thermo Scientific), GFP-trap beads (Proteintech) or Flag-M2 beads (Sigma-Aldrich) (see volumes in **Supplementary Table 8**) were washed twice with 1 mL of PBB buffer (150 mM NaCl, 50 mM Tris-HCl pH 8, 0.5% IGEPAL CA-630, 5 mM MgCl₂, 1 mM DTT and 1x cOmplete protease inhibitor cocktail (Roche)) or PBB-Tx (150 mM NaCl, 50 mM Tris-HCl pH 7.4, 1 mM EDTA, 1% Triton-X and 1x cOmplete protease inhibitor cocktail (Roche)) for Flag-M2 beads. 2 µg (Western blot) or 5 µg (mass spectrometry) of antibody per reaction was added to the pre-washed beads and incubated for 30 min at room temperature on a rotating wheel. Beads were subsequently washed twice with PBB/PBB-Tx buffer and resuspended in 150 µL of PBB/PBB-Tx per immunoprecipitation reaction. 100 µg (Western blot) or 400 µg (mass spectrometry) of protein extract was added to the antibody-bound beads and incubated for 2 h at 4 °C on a rotating wheel. Following immunoprecipitation, beads were washed three times with PBB buffer, then resuspended in 25 µL of 2x Laemmli buffer and boiled at 95 °C for 5 min for elution. For analysis, samples were run on NuPAGE Bis-Tris gels (Thermo Scientific) as detailed in the MS sample preparation and Western Blot sections.

### Western blot

Protein samples for Western blotting were denatured by boiling in 2x Laemmli buffer (Sigma-Aldrich) at 95 °C for 5 min and separated on 4-12% Bis-Tris NuPAGE gels (Thermo Scientific) in 1x MOPS buffer (Thermo Scientific) at 170 V for 60 min. Proteins were transferred to a PVDF membrane (Bio Rad) using a semi-dry blotting system (Power Blotter, Thermo Scientific) for 30 to 60 min at 25 mA. The membrane was blocked for 1 h in PBS containing 0.1% Tween-20 (PBS-T) and 5% skim milk (Nacalai Tesque), then incubated overnight at 4 °C with the primary antibodies (**Supplementary Table 8**). After 3 PBS-T washes, the membrane was incubated for 1 h with horseradish peroxidase-conjugated secondary antibodies (1:5,000 in PBS-T + 5% milk), then washed three more times with PBS-T before detection. For signal detection, Pierce ECL Western blotting substrate (Thermo Scientific) or Amersham ECL prime Western blotting detection reagent (Cytiva) were added to the membrane according to manufacturer’s instructions and imaging was performed using a ChemiDoc MP Imaging System (Bio-Rad). The molecular weight standards used included PageRuler Plus Prestained Protein Ladder (Thermo Scientific) and MagicMark XP Western Protein Standard (Thermo Scientific).

### Chromatin immunoprecipitation (ChIP)

Cells were washed twice with pre-chilled, autoclaved 1x PBS, then cross-linked using 1% formaldehyde (Thermo Fisher Scientific) for 20 min. The reaction was quenched by gently swirling 0.3 M glycine-PBS solution for 5 min. Cross-linked cells were subsequently collected by scraping and pelleted by centrifugation at 1,000 *g* for 5 min. For mass spectrometry applications, a double cross-linking strategy was employed consisting of 2 mM dithiobis(succinimidyl propionate) (DSP) (Thermo Fisher Scientific) for 45 min followed by 1% formaldehyde for 20 min as described previously^36^.

For sonication, cross-linked cell pellets were first lysed in LyB1 buffer (140 mM NaCl, 50 mM Tris-HCl pH 8, 250 mM sucrose, 1 mM EDTA, 10% (v/v) glycerol, 0.5% IGEPAL CA-630, 0.25% TWEEN 20 and 0.25% Triton-X100) for 15 min at 4 °C on a rotating wheel. Cells were then pelleted and subjected to a second lysis step in LyB2 buffer (200 mM NaCl, 10 mM Tris-HCl pH 8, 1 mM EDTA, 0.5 mM EGTA and 1x cOmplete protease inhibitor cocktail) for 10 min at 4 °C on a rotating wheel. Cells were pelleted, resuspended in sonication buffer (50 mM Tris-HCl, 1% (v/v) SDS, 10 mM EDTA and 1x cOmplete protease inhibitor cocktail) at a volume approximately 3 times that of the cell pellet volume. Chromatin was sheared using an EpiShear probe sonicator (Active Motif) for 25 cycles (15 seconds ON, 30 seconds OFF). To verify successful sonication, an aliquot was treated with 100 µg RNase A (Thermo Scientific) for 1 h at 37 °C in 0.2 M NaCl buffer, followed by 50 µg proteinase K digestion (Roche) for 2 h at 62 °C. DNA was purified using the QIAquick PCR purification kit and fragment size was evaluated by agarose gel electrophoresis. For ChIP reactions, 25/720 μg (DNA slot blot/MS) of sonicate was incubated overnight at 4 °C with 0.8/10 μg antibody (**Supplementary Table 8**) diluted in 5.5 volumes of PBB buffer (180 mM NaCl, 50 mM Tris–HCl pH 8, 0.25% IGEPAL CA-630, 5 mM MgCl₂, 1 mM DTT and 1x cOmplete protease inhibitor cocktail). Per ChIP reaction, 12.5/150 μL (DNA slot blot/MS) of Dynabeads Protein G magnetic beads (Thermo Scientific) were washed twice with 2 mL PBB complemented with 15 μg/μL BSA (Sigma-Aldrich), followed by one wash with PBB+BSA supplemented with 3/30 μg (DNA slot blot/MS) of sheared salmon sperm (Thermo Scientific) for 5 min on a rotating wheel. After overnight incubation, each ChIP reaction was mixed with the bead aliquots and was incubated for 2 h at 4 °C. Subsequently, beads were washed 3 or 5 times (for MS and slot blot applications, respectively) with 1 mL PBB (150 mM NaCl, 50 mM Tris–HCl pH 8, 0.5% IGEPAL CA-630, 5 mM MgCl₂, 1 mM DTT and 1x cOmplete protease inhibitor cocktail (Roche)) and once with 1x filtered TE buffer. For MS analyses, beads were resuspended in 25 µL of 2x Laemmli buffer (Sigma-Aldrich) and boiled for 10 min at 95°C then processed as described in the MS sample preparation section. For DNA slot blot analyses, chromatin was eluted twice in 60 μL elution buffer (1% (v/v) SDS and 0.1 M NaHCO3) for 30 min at 65 °C, 1,000 rpm each. Eluted chromatin was then reverse crosslinked by adding 20 mM NaCl and incubated at 65 °C overnight. On the following day, ChIP DNA eluates were subsequently treated with RNAse and proteinase K as described above.

### MS sample preparation, data acquisition and analysis

Immunoprecipitation elutes were separated on a 12% Bis-Tris NuPAGE gel (Thermo Scientific) for 10 min (IP-MS) or 20 min (ChIP-MS) in 1x MOPS buffer (Thermo Scientific) at 170 V. Following electrophoresis, proteins were fixed and stained using the Colloidal Blue Staining Kit (Thermo Scientific) according to the manufacturer’s instructions for approximately 3 h, then destained overnight in HPLC-grade water. For MS sample preparation, each sample lane was divided in 1 (IP-MS) or 2 equal fractions (ChIP-MS) and each fraction was cut into approximately 1x1 mm pieces. Gel pieces were destained twice with 50 mM ammonium bicarbonate in 50% EtOH, then dehydrated twice in 100% acetonitrile solution and, to ensure complete removal of acetonitrile, dried in a vacuum centrifuge for 5 min. Next, disulfide bonds were broken by incubating the gel pieces in reduction buffer (50 mM ammonium bicarbonate and 10 mM DTT) for 1 h at 56 °C, followed by 45 min incubation in alkylation buffer (50 mM ammonium bicarbonate and 55 mM iodoacetamide (Sigma-Aldrich)) at room temperature and in the dark. Subsequently, gel pieces were washed once in 50 mM ammonium bicarbonate solution and once in 100% acetonitrile. The gel pieces were rehydrated and dehydrated once more, followed by a final dehydration in acetonitrile, and subsequent drying in a vacuum centrifuge (Concentrator Plus (Eppendorf)) for 5 min. Next, proteins were digested overnight at 37 °C using 2 μg sequencing-grade trypsin (Promega) in 50 mM ammonium bicarbonate. After overnight digestion, the supernatant was collected, and the digested peptides left in the gel pieces were extracted for 15 min with extraction buffer (50 mM ammonium bicarbonate, 30% acetonitrile and 3% trifluoroacetic acid) followed by a dehydration step (acetonitrile).

These two steps were repeated once more and a final dehydration step was included. For each step, supernatants were pooled and subsequently concentrated using a Concentrator Plus (Eppendorf) for 2-3 h to a final volume appropriate for stage-tipping (< 200 μL). For stage-tipping, two layers of C18 extraction disks (3M) were placed into 200 μL pipette tips and activated with 50 μL methanol. After activation, the material was equilibrated using 50 μL solution B (80% acetonitrile, 0.5% formic acid), rinsed twice with 50 μL solution A (0.5% formic acid) and then loaded with the peptide samples, followed by a final wash with 50 μL solution A.

Peptides were analyzed using nanoflow liquid chromatography on an EASY-nLC 1200 system coupled to a Q Exactive HF mass spectrometer (Thermo Scientific). Separation was performed on a 75 μm inner diameter in-house packed C18 reversed-phase column, fitted with an integrated 8 μm emitter tip and packed with ReproSil-Pur C18-AQ 1.9 μm resin (Dr. Maisch). The column was mounted on an Easy Flex Nano Source and maintained at 40 °C using a column oven (Sonation). Peptide elution was carried out with an 88-min gradient ranging from 2% to 40% solution B (80 % acetonitrile, 0.1 % formic acid) at a constant flow rate of 225 nL/min. The spray voltage was set to 2.2 kV. The mass spectrometer was operated in data-dependent acquisition mode, collecting one full MS scan followed by up to 20 MS/MS spectra (Top20 method). Full MS scans were acquired at a resolution of 60,000 with a maximum injection time of 20 ms, and MS/MS scans at a resolution of 15,000 with a maximum injection time of 75 ms.

Raw data were processed using MaxQuant^53^ (v1.5.2.8) with preset standard settings for label-free quantitation using at least 2 LFQ ratio counts based on the MaxLFQ algorithm^54^. Carbamidomethylation was set as a fixed modification while acetyl, methyl, methionine oxidation and protein N-acetylation were considered as variable modifications. Peptide and protein FDR were enforced at 0.01 with match between runs option activated. Missing values were imputed and proteins not quantified in at least 4 replicates were excluded from the analysis using an in-house Python script, which was also used to generate volcano plots. For the cross-link analysis MaxQuant (v2.6.4.0) was used and the default settings were modified as follows: maximum missed cleavages to 3, cross-linking type to MS2-cleavable, cross-linker to DSBU and minimum peptide length to 5. This analysis was run on two experimental setups: digestion with trypsin alone and digestion with trypsin and LysC.

### DNA slot blot

Before blotting, purified DNA recovered by ChIP was denatured in 1x denaturation buffer (0.8 M NaOH and 20 mM EDTA, pH 8) at 100 °C for 10 min and neutralized with the addition of ice-cold 1 M ammonium acetate. DNA samples were then spotted on a positively charged nylon membrane (Hybond, N+, Amersham) pre-washed with autoclaved Milli-Q water and 6x SSC. Next, the membrane was completely air-dried, and the blotted DNA was UV crosslinked twice at 120 mJ using a Stratalinker UV Crosslinker (Stratagene). Telomeric DNA was detected using the TeloTTAGG telomere length assay kit (Roche) as described in Southern blot analysis of Telomere Restriction Fragment length method.

### Nucleosome preparation

Telomeric DNA templates containing 10 repeats of 157 bp telomeric DNA separated by AvaI sites were prepared employing established protocols^55^. Briefly, the plasmid containing telomeric DNA template was amplified in SURE2 *E. coli* strain in terrific broth at 30°C. The cells were grown for ∼18 h and plasmids were purified by alkaline lysis. The concentration of purified plasmid was adjusted to 2 mg/mL followed by overnight digestion at 37°C with AvaI at an enzyme concentration of 100 units/mg of plasmid. The digested plasmid was fractionated on a Resource Q column employing a linear gradient of buffer containing lithium chloride (20 mM Tris, 300-750 mM LiCl, 1 mM EDTA) and further polished by MonoQ column to obtain 157 bp telomeric DNA templates^56^. Individual histones were overexpressed in *E. coli* and purified as previously described. The histone octamer was reconstituted by mixing histones H2A, H2B, H3 and H4 under denaturing conditions in stoichiometric ratio of 1.2:1.2:1:1 and dialysing against high salt. This histone octamer was further purified by size exclusion chromatography and employed for nucleosome reconstitution.

Nucleosome reconstitution was performed via salt gradient dialysis utilizing 10 kDa MWCO Slide-A-Lyzer (Thermo Fisher Singapore). Reaction mixtures were assembled by sequentially adding autoclaved water, 1 M Tris (pH 7.5), 4 M LiCl, 0.5 M EDTA, 0.5 M DTT, human histone octamer (hHO), and DNA. The final composition of the assembly mixture was 20 mM Tris (pH 7.5), 2 M LiCl, 1 mM EDTA, 1 mM DTT, 4.8–7.2 μM hHO, and 6 μM DNA.

Initially, the samples were equilibrated against a high-salt buffer (20 mM Tris pH 7.5, 2 M LiCl, 1 mM EDTA, 1 mM DTT) for 30 minutes at room temperature. Subsequently, salt concentration was lowered using a continuous-flow system in which low-salt buffer (20 mM Tris pH 7.5, 1 mM EDTA, 1 mM DTT) was introduced while an equivalent volume of the buffer mixture was removed. For small-scale preparations, this involved pumping 500 mL of low-salt buffer into a reservoir containing 200 mL of high-salt buffer, concluding with a 4-hour dialysis step in pure low-salt buffer.

### Electrophoretic mobility assay (EMSA)

To analyse protein-nucleosome interactions, HOT1 was titrated against reconstituted telomeric mononucleosomes and dinucleosomes at molar ratios of 0:1, 2:1, and 4:1 (protein:nucleosome). Prior to the assay, HOT1 was buffer-exchanged into 1x physiological buffer (20 mM HEPES pH 7.5, 130 mM KCl, 20 mM NaCl, 1 mM DTT, 0.1 mM EDTA, 1 mM MgCl2, 5% glycerol) using Amicon Ultra 0.5 mL centrifugal filters (Merck Millipore). The reconstituted nucleosomes, HOT1, DYNLL1 and buffer components were mixed in a PCR tube to obtain a final concentration of 20 mM HEPES (pH 7.5), 100 ng/ul of DNA (nucleosome), 100 mM KCl, 1 mM MgCl2, 0.1 mM EDTA and 1 mM DTT. Reactions were mixed by gentle tapping and a brief spin, followed by incubation on ice for 20 min. Subsequently, 6x TriTrack DNA loading dye was added, and samples (∼400 ng DNA per well) were resolved on a 0.80% agarose gel in 0.25x TBE. Electrophoresis was performed in ice-cold 0.25x TBE buffer at 110 V for 71 min.

### MgCl2 precipitation assay

The reconstituted nucleosomes, HOT1, DYNLL1 and buffer components were mixed in a PCR tube to obtain a final concentration of 20 mM HEPES (pH 7.5), 100 ng/ul of DNA (nucleosome), 100 mM KCl, 1 mM MgCl2, 0.1 mM EDTA and 1 mM DTT. Reactions were mixed by gentle tapping and a brief spin, followed by incubation on ice for 20 min. To induce nucleosome precipitation, MgCl₂ was added to a final concentration of 20 mM, followed by a second incubation on ice for 20 min. Insoluble complexes were pelleted by centrifugation at 20,000 g for 20 min. The supernatant was carefully recovered, and 5x SDS loading dye was added. The pellet was resuspended in 1x Tris-Glycine buffer mixed with 5x SDS loading dye. Both fractions were denatured at 95°C for 5 min and resolved on 15% SDS-PAGE gels.

### Cryo-EM sample preparation

NCP-HOT1 and NCP-HOT1-DYNLL1 were generated with a molar stoichiometry of 1:1 as described above at a telomeric NCP concentration of 1 mg/ml. The samples were dialysed overnight against 20 mM HEPES (pH 7.5), 20 mM KCl, 1 mM DTT, 1 mM EDTA using a Slide-A-dialyzer. The samples were recovered and centrifuged at 20,000 g for 20 min. 4 μL of the supernatant were each applied to a glow-discharged Quantifoil R1.2/1.3 Au grid and blotted for 1.5 s with a blot force of 1 at 4 °C and relative humidity of 100% and flash frozen in liquid ethane at -180 °C. The grids were imaged using an ARCTICA cryo-electron microscope operating at 200 kV.

## Data availability

The mass spectrometry data have been deposited to the ProteomeXchange Consortium via the PRIDE partner repository^57^ with the dataset identifier PXD076300.

## Acknowledgments

We are grateful to all members of the Kappei lab for advice and discussions. This research was supported by the National Research Foundation Singapore and the Singapore Ministry of Education under its Research Centres of Excellence initiative [DK], an NMRC Open Fund Young Investigator Research Grant (MOH-001522) [AS] and a NTU Presidential Postdoctoral Fellowship (023462-00001) [AS]. The authors thank the Spatial Biology Core (SBC) and the Quantitative Proteomics Core (QPC) at the Cancer Science Institute of Singapore (CSI) for their support, the NTU Institute of Structural Biology for cryo-EM usage and the Protein Production Platform (PPP) at the NTU School of Biological Sciences for the DYNLL1 purification.

## Author contributions

AH and DK conceived the study and designed experiments. AH performed and analyzed most experiments with help from NZA, VK, MY, JHT, CYG, FB and DK. NK, LN and AS performed and analysed the nucleosome binding tests and cryoEM screening. AH and DK analysed the data and wrote the manuscript with input from all authors.

---

**Supplementary Figure 1.**
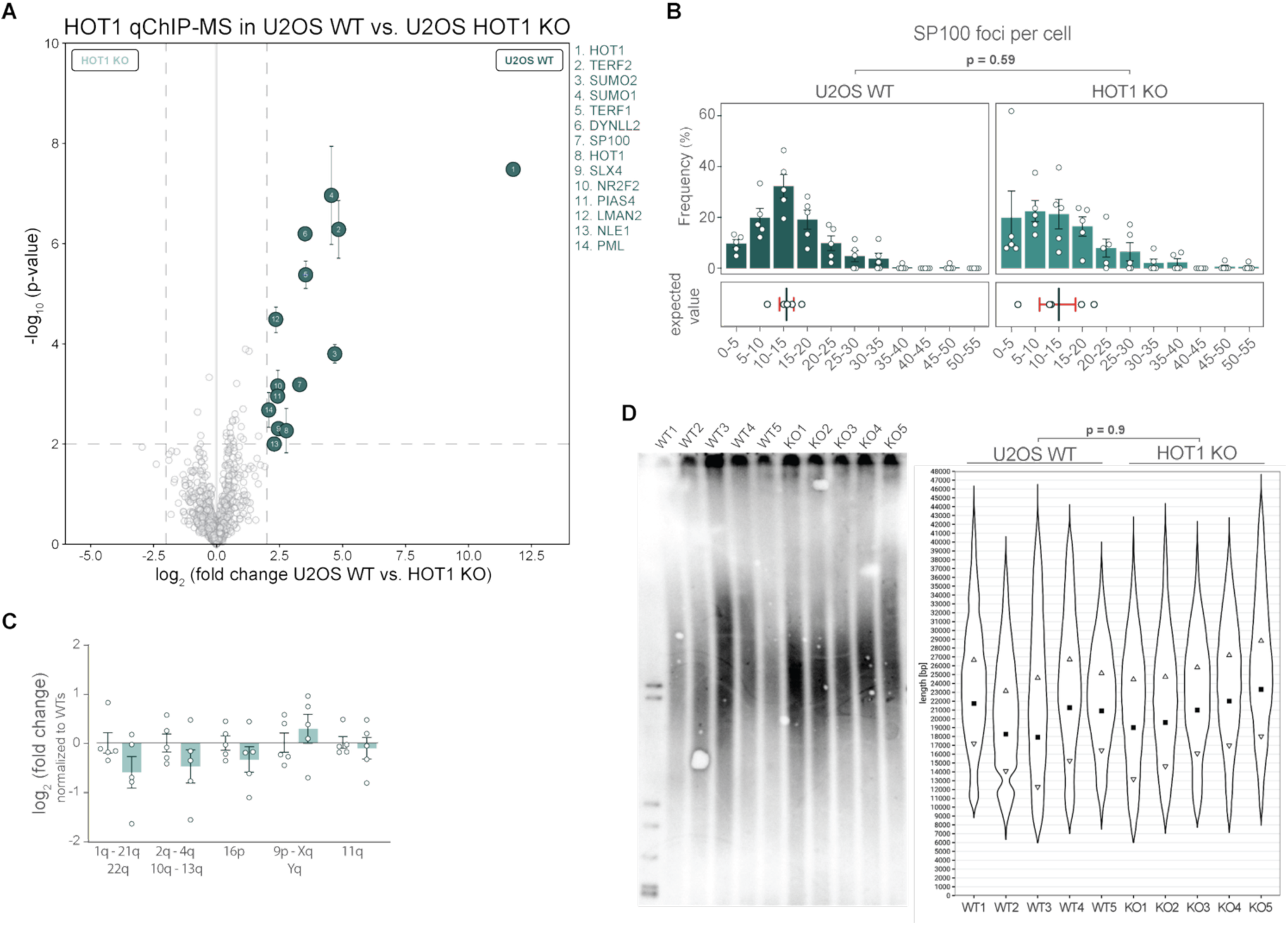
HOT1 associates with ALT-positive telomeres but does not affect APB assembly, TERRA levels or telomere length in U2OS cells. **(A)** Volcano plot comparing HOT1 qChIP-MS from 5 U2OS WT clones vs. 5 U2OS HOT1 KO clones. Enriched proteins are distinguished from background binders by a two-dimensional cut-off of > 4-fold enrichment and p-value < 0.01 (*n=5*). **(B)** Quantification of the number of PML bodies per cell in U2OS WT clones (*n=5, 39-52 nuclei analysed per clone*) and U2OS HOT1 KO clones (*n=5, 32-66 nuclei analysed per clone*). Upper plot: the frequency of cells for the indicated number of PML bodies, error bars represent SEM across clones. Lower plot: the vertical line represents the fitted expected number of PML bodies (cumulative link mixed model (CLMM) with a proportional-odds logit link), the error bar represents the 95% confidence interval. p-value is calculated using likelihood ratio test. **(C)** RT-qPCR analyses of TERRA expression from the indicated chromosome arms in each 5 U2OS WT and HOT1 KO clones (*n=5*). Each clone (represented by dots) was quantified using the average of technical triplicates. Mean values are shown with error bars representing SD and p-values were calculated for the biological replicates using Welch’s t-test. **(D)** Telomere Restriction Fragment (TRF) analysis of each 5 U2OS WT and HOT1 KO clones (*n=5*) using pulsed-field gel electrophoresis (PFGE) (left). Average telomere length for each clone was calculated and plotted using WALTER^52^ (right). The violin plot indicates the variation in the intensity of the telomere smear. The weighted median is indicated by a solid square and the 1^st^ and 3^rd^ quartile by the arrow heads. The significance is tested by Tamhane-Dunnett test.

**Supplementary Figure 2.**
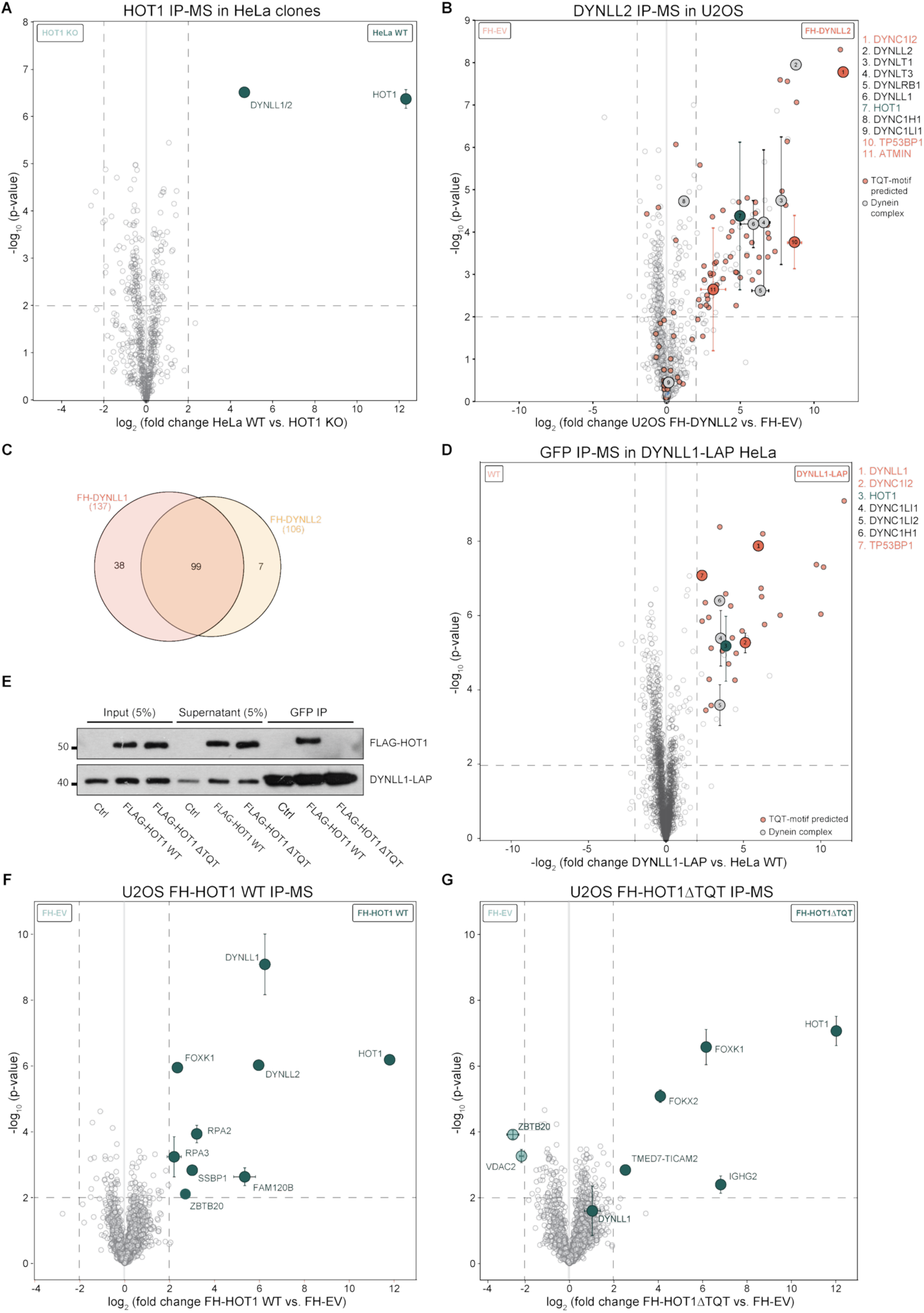
HOT1 interacts with DYNLL1/2 via its TQT motif. **(A)** Volcano plot comparing HOT1 IP-MS reactions from 5 HeLa WT clones and 5 HeLa HOT1 KO clones (*n=5*). **(B)** Volcano plot comparing FLAG IP-MS reactions from U2OS cells expressing either FLAG-DYNLL2 or FLAG empty vector (EV) (*n=4*). Proteins predicted to interact with DYNLL2 are highlighted in red and members of the dynein complex are highlighted in grey. **(C)** Venn-diagram of enriched proteins in FH-DYNLL1 and FH-DYNLL2 IP-MS data. **(D)** Volcano plot comparing GFP IP-MS reactions in HeLa DYNLL1-LAP tagged^40^ (DYNLL1 is fused to GFP) and HeLa WT (*n=4*). Enriched proteins are distinguished from background as indicated in **(B)**. **(E)** Immunoprecipitation of DYNLL1-LAP performed in HeLa cell expressing either FLAG–HOT1 WT or FLAG-HOT1ΔTQT. **(F)** Volcano plot comparing FLAG IP-MS reactions from 5 U2OS HOT1 KO clones expressing either FLAG-HOT1 WT or FLAG-EV (*n=5*). **(G)** Volcano plot comparing FLAG IP-MS reactions from 5 U2OS HOT1 KO clones expressing either FLAG-HOT1ΔTQT or FLAG-EV (*n=5*). All pull-downs were analysed by label-free quantification. Specifically enriched proteins are distinguished from background binders by a two-dimensional cut-off of > 4-fold enrichment and p-value < 0.01. Two-dimensional error bars represent the standard deviation based on iterative imputation cycles during the label-free analysis to substitute missing values (e.g. no detection in the HOT1 KO control).

**Supplementary Figure 3.**
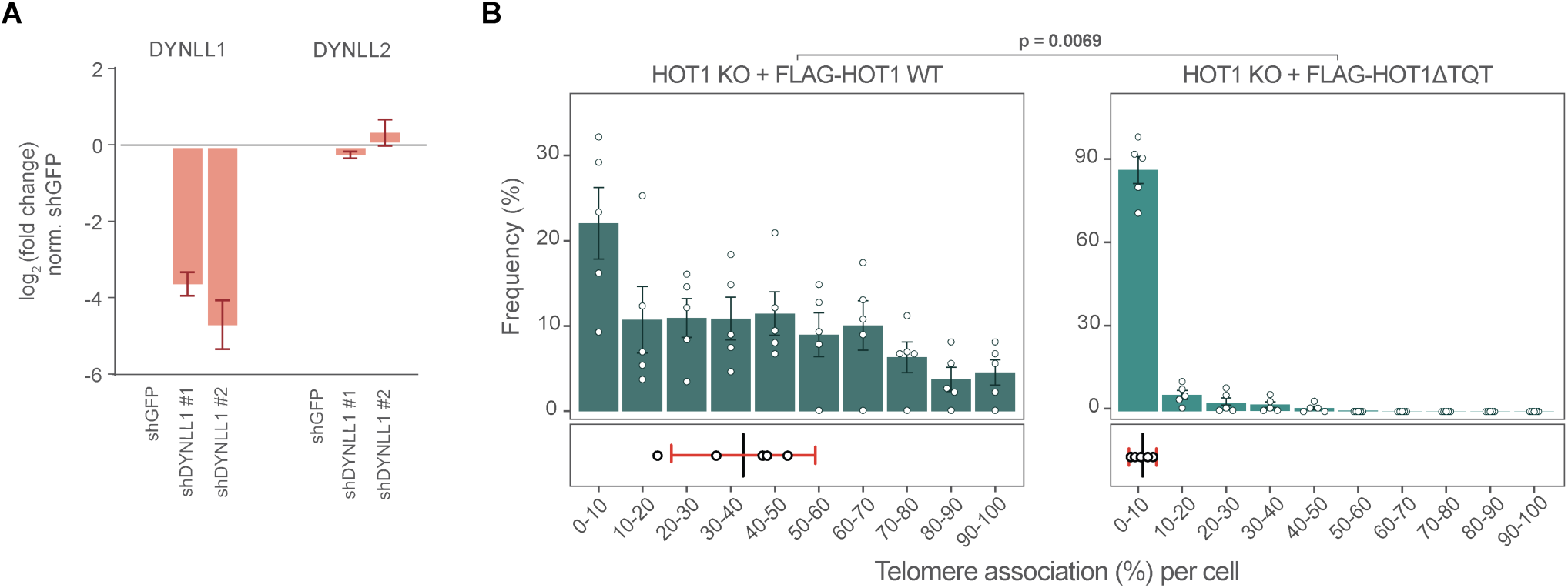
DYNLL1/2-mediated HOT1 dimerization is required for HOT1-telomere localization *in vivo*. **(A)** DYNLL1/2 mRNA expression in U2OS cells upon shGFP, shDYNLL1 #1 or shDYNLL1 #2 knock-down. **(B)** Quantification of the HOT1 percent telomere association per cell in U2OS KO + FH-HOT1 WT clones and U2OS KO + FH-HOT1ΔTQT clones (*n=5, 74-107 nuclei analysed per clone*). Upper plot: the frequency of cells for the indicated percentage of telomere association with HOT1, error bars represent the SEM across clones. Lower plot: the vertical line represents the fitted expected percentage of telomere association (zero-inflated beta GLMM), the error bar represents the 95% confidence interval. p-value was calculated using likelihood ratio test.

**Supplementary Figure 4.**
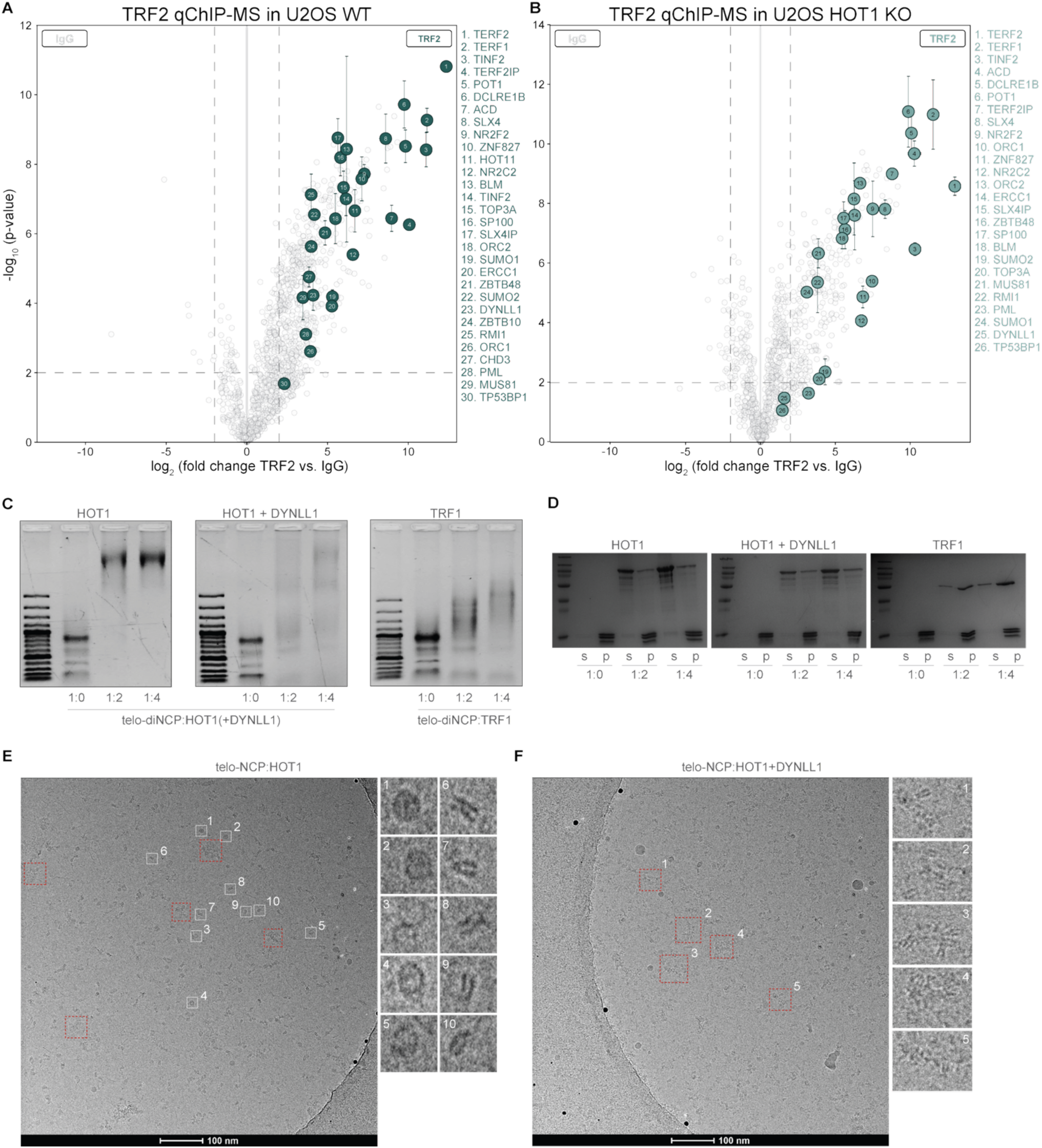
HOT1 binds telomeric nucleosomes. **(A-B)** Volcano plot comparing TRF2 vs. IgG qChIP-MS reactions in 5 U2OS WT clones (*n=5*) **(A)** and 5 U2OS HOT1 KO clones (*n=5*) **(B)**. Specifically enriched proteins are distinguished from background by a two-dimensional cut-off with fold enrichment > 4 and p-value < 0.01. Two-dimensional error bars represent the standard deviation based on iterative imputation cycles during the label-free analysis to substitute missing values (e.g. no detection in the IgG control). (**C**) Electrophoretic mobility assay of telomeric di-nucleosomes bound with HOT1 (left), HOT1+DYNLL1 (middle) and TRF1 (right). Numbers indicate the ratios of NCP over the purified proteins. (**D**) Electrophoresis analysis following MgCl2 precipitation of mono-nucleosomes mixed to HOT1, HOT1+DYNLL1 and TRF1. s indicates supernatant loading and p indicates pellet loading. Numbers indicate the ratios of NCP over the purified proteins. (**E-F**) Cryo-EM screening of (**E**) HOT1 and (**F**) HOT1+DYNLL1 interaction with telomeric mono-nucleosomes. Cryo-EM visualizations showed nucleosomes (white squares) along with aggregates (red squares) bound by HOT1 and HOT1+DYNLL1 binding to telomeric nucleosomes. Representative views are shown.

## Supplementary Tables

**Supplementary Table 1 | U2OS HOT1 qChIP-MS data**

**Supplementary Table 2 | U2OS & HeLa WT vs KO HOT1 IP-MS data**

**Supplementary Table 3 | U2OS & HeLa DYNLL1/2 IP-MS data**

**Supplementary Table 4 | U2OS HOT1 KO FH-HOT1 IP-MS data**

**Supplementary Table 5 | U2OS HOT1 WT vs KO TRF2 qChIP-MS data**

**Supplementary Table 6 | HOT1 nucleosomes XL-MS data**

**Supplementary Table 7 |Oligonucleotides used in this study**

**Supplementary Table 8 |Antibodies used in this study**

## References

1. Palm, W. & Lange, T. de. How Shelterin Protects Mammalian Telomeres. Annu. Rev. Genet. 42, 301–334 (2008).

2. Lange, T. de. How Telomeres Solve the End-Protection Problem. Science 326, 948–952 (2009).

3. Levy, M. Z., Allsopp, R. C., Futcher, A. B., Greider, C. W. & Harley, C. B. Telomere end-replication problem and cell aging. J. Mol. Biol. 225, 951–960 (1992).

4. Kim, N. W. et al. Specific Association of Human Telomerase Activity with Immortal Cells and Cancer. Science 266, 2011–2015 (1994).

5. Bryan, T. M., Englezou, A., Dalla-Pozza, L., Dunham, M. A. & Reddel, R. R. Evidence for an alternative mechanism for maintaining telomere length in human tumors and tumor-derived cell lines. Nat. Med. 3, 1271– 1274 (1997).

6. Henson, J. D. & Reddel, R. R. Assaying and investigating Alternative Lengthening of Telomeres activity in human cells and cancers. FEBS Lett. 584, 3800–3811 (2010).

7. Heaphy, C. M. et al. Prevalence of the Alternative Lengthening of Telomeres Telomere Maintenance Mechanism in Human Cancer Subtypes. Am. J. Pathol. 179, 1608–1615 (2011).

8. Zhang, J.-M., Yadav, T., Ouyang, J., Lan, L. & Zou, L. Alternative Lengthening of Telomeres through Two Distinct Break-Induced Replication Pathways. Cell Rep. 26, 955–968.e3 (2019).

9. Yeager, T. R. et al. Telomerase-negative immortalized human cells contain a novel type of promyelocytic leukemia (PML) body. Cancer Res. 59, 4175–9 (1999).

10. Draskovic, I. et al. Probing PML body function in ALT cells reveals spatiotemporal requirements for telomere recombination. Proc. Natl. Acad. Sci. 106, 15726–15731 (2009).

11. Chung, I., Leonhardt, H. & Rippe, K. De novo assembly of a PML nuclear subcompartment occurs through multiple pathways and induces telomere elongation. J. Cell Sci. 124, 3603–3618 (2011).

12. Jiang, W.-Q., Nguyen, A., Cao, Y., Chang, A. C.-M. & Reddel, R. R. HP1-Mediated Formation of Alternative Lengthening of Telomeres-Associated PML Bodies Requires HIRA but Not ASF1a. PLoS ONE 6, e17036 (2011).

13. Gauchier, M. et al. SETDB1-dependent heterochromatin stimulates alternative lengthening of telomeres. Sci. Adv. 5, eaav3673 (2019).

14. Zhang, H. et al. Nuclear body phase separation drives telomere clustering in ALT cancer cells. Mol. Biol. Cell 31, 2048–2056 (2020).

15. Bryan, T. M., Englezou, A., Gupta, J., Bacchetti, S. & Reddel, R. R. Telomere elongation in immortal human cells without detectable telomerase activity. EMBO J. 14, 4240–4248 (1995).

16. Londoño-Vallejo, J. A., Der-Sarkissian, H., Cazes, L., Bacchetti, S. & Reddel, R. R. Alternative Lengthening of Telomeres Is Characterized by High Rates of Telomeric Exchange. Cancer Res. 64, 2324–2327 (2004).

17. Takai, H., Smogorzewska, A. & Lange, T. de. DNA Damage Foci at Dysfunctional Telomeres. Curr. Biol. 13, 1549–1556 (2003).

18. Ng, L. J., Cropley, J. E., Pickett, H. A., Reddel, R. R. & Suter, C. M. Telomerase activity is associated with an increase in DNA methylation at the proximal subtelomere and a reduction in telomeric transcription. Nucleic Acids Res. 37, 1152–1159 (2009).

19. Episkopou, H. et al. Alternative Lengthening of Telomeres is characterized by reduced compaction of telomeric chromatin. Nucleic Acids Res. 42, 4391–4405 (2014).

20. Cesare, A. J. & Griffith, J. D. Telomeric DNA in ALT Cells Is Characterized by Free Telomeric Circles and Heterogeneous t-Loops. Mol. Cell. Biol. 24, 9948–9957 (2004).

21. Henson, J. D. et al. DNA C-circles are specific and quantifiable markers of alternative-lengthening-of-telomeres activity. Nat. Biotechnol. 27, 1181–1185 (2009).

22. Heaphy, C. M. et al. Altered Telomeres in Tumors with ATRX and DAXX Mutations. Science 333, 425–425 (2011).

23. Napier, C. E. et al. ATRX represses alternative lengthening of telomeres. Oncotarget 6, 16543–16558 (2015).

24. Li, F. et al. ATRX loss induces telomere dysfunction and necessitates induction of alternative lengthening of telomeres during human cell immortalization. EMBO J. 38, EMBJ201796659 (2019).

25. Turkalo, T. K. et al. A non-genetic switch triggers alternative telomere lengthening and cellular immortalization in ATRX deficient cells. Nat. Commun. 14, 939 (2023).

26. Kappei, D. et al. Phylointeractomics reconstructs functional evolution of protein binding. Nat. Commun. 8, 14334 (2017).

27. Braun, H. et al. ZNF524 directly interacts with telomeric DNA and supports telomere integrity. Nat. Commun. 14, 8252 (2023).

28. Conomos, D., Reddel, R. R. & Pickett, H. A. NuRD–ZNF827 recruitment to telomeres creates a molecular scaffold for homologous recombination. Nat. Struct. Mol. Biol. 21, 760–770 (2014).

29. Déjardin, J. & Kingston, R. E. Purification of Proteins Associated with Specific Genomic Loci. Cell 136, 175– 186 (2009).

30. Conomos, D. et al. Variant repeats are interspersed throughout the telomeres and recruit nuclear receptors in ALT cells. J. Cell Biol. 199, 893–906 (2012).

31. Bluhm, A. et al. ZBTB10 binds the telomeric variant repeat TTGGGG and interacts with TRF2. Nucleic Acids Res. 47, 1896–1907 (2019).

32. Jahn, A. et al. ZBTB48 is both a vertebrate telomere-binding protein and a transcriptional activator. EMBO Rep. 18, 929–946 (2017).

33. Li, J. S. Z. et al. TZAP: A telomere-associated protein involved in telomere length control. Science 355, 638– 641 (2017).

34. Kappei, D. et al. HOT1 is a mammalian direct telomere repeat-binding protein contributing to telomerase recruitment. EMBO J. 32, 1681–1701 (2013).

35. Feng, X. et al. The telomere-associated homeobox-containing protein TAH1/HMBOX1 participates in telomere maintenance in ALT cells. J. Cell Sci. 126, 3982–3989 (2013).

36. Yong, W. K. et al. qChIP-MS reveals the local chromatin composition by label-free quantitative proteomics. Nat. Commun. 17, 6857 (2026).

37. Mok, Y.-K., Lo, K. W.-H. & Zhang, M. Structure of Tctex-1 and Its Interaction with Cytoplasmic Dynein Intermediate Chain*. J. Biol. Chem. 276, 14067–14074 (2001).

38. Becker, J. R. et al. The ASCIZ-DYNLL1 axis promotes 53BP1-dependent non-homologous end joining and PARP inhibitor sensitivity. Nat. Commun. 9, 5406 (2018).

39. Howe, J. et al. LC8 enhances 53BP1 foci through heterogeneous bridging of 53BP1 oligomers. eLife 14, e102179 (2025).

40. Hubner, N. C. et al. Quantitative proteomics combined with BAC TransgeneOmics reveals in vivo protein interactions. J. Cell Biol. 189, 739–754 (2010).

41. Barbar, E. Dynein Light Chain LC8 Is a Dimerization Hub Essential in Diverse Protein Networks †. Biochemistry 47, 503–508 (2008).

42. Benison, G., Karplus, P. A. & Barbar, E. Structure and Dynamics of LC8 Complexes with KXTQT-Motif Peptides: Swallow and Dynein Intermediate Chain Compete for a Common Site. J. Mol. Biol. 371, 457–468 (2007).

43. Jespersen, N. et al. Systematic identification of recognition motifs for the hub protein LC8. Life Sci. Alliance 2, e201900366 (2019).

44. Court, R., Chapman, L., Fairall, L. & Rhodes, D. How the human telomeric proteins TRF1 and TRF2 recognize telomeric DNA: a view from high-resolution crystal structures. EMBO Rep. 6, 39–45 (2005).

45. Fairall, L., Chapman, L., Moss, H., Lange, T. de & Rhodes, D. Structure of the TRFH Dimerization Domain of the Human Telomeric Proteins TRF1 and TRF2. Mol. Cell 8, 351–361 (2001).

46. Rose, R. B., Endrizzi, J. A., Cronk, J. D., Holton, J. & Alber, T. High-Resolution Structure of the HNF-1α Dimerization Domain †, ‡. Biochemistry 39, 15062–15070 (2000).

47. Lee, J. H. et al. Regulation of telomere homeostasis and genomic stability in cancer by N6-adenosine methylation (m6A). Sci. Adv. 7, eabg7073 (2021).

48. Price, A. J., Manjegowda, M. C., Kain, J., Anandh, S. & Bochkis, I. M. Hdac3, Setdb1, and Kap1 mark H3K9me3/H3K14ac bivalent regions in young and aged liver. Aging Cell 19, e13092 (2020).

49. Denslow, S. A. & Wade, P. A. The human Mi-2/NuRD complex and gene regulation. Oncogene 26, 5433– 5438 (2007).

50. Hossain, M. & Stillman, B. Opposing roles for DNA replication initiator proteins ORC1 and CDC6 in control of Cyclin E gene transcription. eLife 5, e12785 (2016).

51. Deng, Z., Norseen, J., Wiedmer, A., Riethman, H. & Lieberman, P. M. TERRA RNA Binding to TRF2 Facilitates Heterochromatin Formation and ORC Recruitment at Telomeres. Mol. Cell 35, 403–413 (2009).

52. Lyčka, M. et al. WALTER: an easy way to online evaluate telomere lengths from terminal restriction fragment analysis. BMC Bioinform. 22, 145 (2021).

53. Cox, J. & Mann, M. MaxQuant enables high peptide identification rates, individualized p.p.b.-range mass accuracies and proteome-wide protein quantification. Nat. Biotechnol. 26, 1367–1372 (2008).

54. Cox, J. et al. Accurate Proteome-wide Label-free Quantification by Delayed Normalization and Maximal Peptide Ratio Extraction, Termed MaxLFQ*. Mol. Cell. Proteom. 13, 2513–2526 (2014).

55. Wong, S. Y. et al. The shelterin component TRF2 mediates columnar stacking of human telomeric chromatin. EMBO J. 43, 87–111 (2024).

56. Soman, A. et al. The human telomeric nucleosome displays distinct structural and dynamic properties. Nucleic Acids Res. 48, 5383–5396 (2020).

57. Perez-Riverol, Y. et al. The PRIDE database at 20 years: 2025 update. Nucleic Acids Res. 53, D543–D553 (2024).

